# Prex1 controls glucose homeostasis by limiting glucose uptake and mitochondrial metabolism in liver through GEF-independent regulation of Gpr21

**DOI:** 10.1101/2025.04.14.648781

**Authors:** Julia Y. Chu, Elpida Tsonou, Polly A. Machin, Kirsty MacLellan-Gibson, Anna Roberts, Stephen A. Chetwynd, Adam T. McCormack, John C. Stephens, Elisa Benetti, Gemma K. Kinsella, David Baker, David C. Hornigold, Heidi C. E. Welch

## Abstract

We investigated the roles of Rac guanine-nucleotide factor (Rac-GEF) Prex1 in glucose homeostasis using *Prex1^−/−^* and catalytically-inactive *Prex1^GD^* mice. Prex1 maintains fasting blood glucose levels and insulin sensitivity through its Rac-GEF activity but limits glucose clearance independently of its catalytic activity, throughout ageing. *Prex1^−/−^*mice on high-fat diet are protected from developing diabetes. The increased glucose clearance in *Prex1^−/−^* mice stems from constitutively enhanced hepatic glucose uptake. Prex1 limits Glut2 surface levels, mitochondrial membrane potential and mitochondrial ATP production, and controls mitochondrial morphology in hepatocytes, independently of its catalytic activity. Prex1 limits GPCR trafficking through an adaptor function, and we identify here the inhibitory orphan GPCR Gpr21 as a Prex1 target. The Gpr21-mediated blockade of glucose uptake and mitochondrial ATP production in hepatocytes requires Prex1. We propose that Prex1 limits glucose clearance by maintaining Gpr21 at the hepatocyte surface, thus limiting hepatic glucose uptake and metabolism.

**Graphical abstract:** 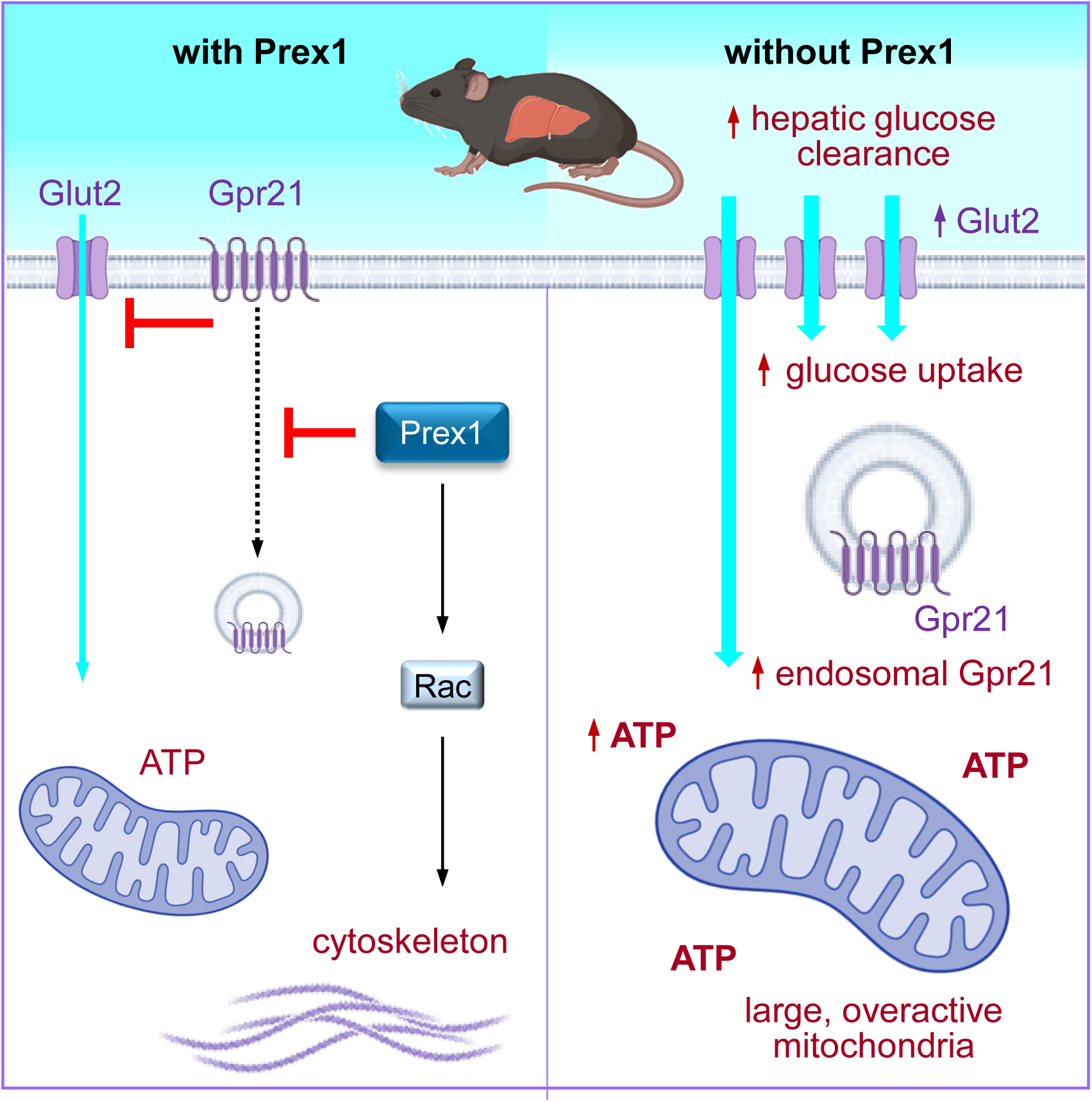

## Introduction

Glucose homeostasis, the maintenance of blood glucose levels within their physiological range, is a vital process. When blood glucose levels rise following a meal, glucose is cleared into the brain and liver independently of insulin, and into adipose and skeletal muscle tissues in an insulin-dependent manner. In the fasting state, blood glucose levels are maintained by the liver, through glycogenolysis and gluconeogenesis. Hyper- and hypoglycaemia can cause symptoms ranging from headache and tiredness to coma and death. Chronic hyperglycaemia leads to metabolic syndrome, comprising obesity, insulin resistance, and type 2 diabetes, with increased risk of nephropathy, neuropathy, cardiovascular disease and cancer (Gerich, 2000). These diseases are rampant in the developing and developed world, and compliance with healthy lifestyle choices is proving insufficient to curb the ever-increasing trends. Medical interventions have limitations, meaning new therapeutic opportunities are needed.

Rac-GTPases, and the guanine-nucleotide exchange factors (GEFs) that activate them, are central controllers of cytoskeletal dynamics, and thereby of cell morphology, adhesion, migration, and secretory processes (Cherfils and Zeghouf, 2013; Rossman et al., 2005). Recently, these proteins are becoming recognised as regulators of glucose homeostasis (Machin et al., 2021; Moller et al., 2019). In skeletal muscle and adipose tissue, they mediate glucose uptake by controlling the insulin-stimulated translocation of the glucose transporter GLUT4 to the plasma membrane, and in pancreatic β cells, they control glucose-stimulated insulin release. Mice with an L332A mutation in the catalytic DH domain of the Rac-GEF Vav2 that reduces catalytic activity show low skeletal muscle mass, hyperglycaemia, glucose intolerance, increased adiposity of brown and white adipose tissues (BAT and WAT), and liver steatosis (Lorenzo-Martin et al., 2020). In contrast, mice with catalytically hyperactive, N-terminally truncated Vav2 show increased muscle mass, dependent on the Rac1-GEF activity, and are resistant to developing metabolic syndrome even when fed a high-fat diet (HFD) (Lorenzo-Martin et al., 2020; Rodriguez-Fdez et al., 2021; Rodriguez-Fdez et al., 2020). Mice deficient in the related Rac-GEF Vav3 show different metabolic phenotypes depending on diet. On normal chow diet, *Vav3^−/−^* mice develop metabolic syndrome, with non-alcoholic fatty liver disease (NAFLD) and type 2 diabetes, whereas on high-fat diet, they are protected from obesity and metabolic syndrome due to increased energy consumption and brown adipose tissue thermogenesis, ultimately through chronic sympatho-excitation enhancing white-to-brown adipose trans-differentiation (Menacho-Marquez et al., 2013). Mice deficient in the Rac-GEF Prex2 show reduced glucose tolerance and cannot sustain insulin-stimulated glucose clearance (Hodakoski et al., 2014). P-Rex2 protein levels are reduced and the activity of the tumour suppressor PTEN is increased in adipose tissue of insulin-resistant humans (Hodakoski et al., 2014). P-Rex2 inhibits PTEN through an adaptor function, leading to elevated levels of the lipid second messenger PIP_3_ generated by phosphoinositide 3-kinase (PI3K), and it was proposed that P-Rex2 controls glucose homeostasis through PTEN inhibition rather than its catalytic Rac-GEF activity (Fine et al., 2009).

P-Rex1 is a widely expressed GEF for the Rac-family small GTPases, which together with P-Rex2 makes up the P-Rex family (Hornigold K, 2018; Welch, 2015; Welch et al., 2002). P-Rex1 and P-Rex2 have a unique synergistic mode of activation by PIP_3_ and the Gβγ subunits of heterotrimeric G proteins, which makes these GEFs ideal mediators of G protein-coupled receptor (GPCR) signalling (Hornigold K, 2018; Welch, 2015; Welch et al., 2002). P-Rex1 is best-known for its roles in the immune and nervous systems, where it is required for a range of pro-inflammatory and immune functions, synaptic plasticity, fine motor skills and social recognition (Hornigold K, 2018; Welch, 2015; Welch et al., 2002), but roles in metabolic processes have also been emerging (Machin et al., 2021). Single nucleotide polymorphisms in the perigenic region of human *PREX1* are correlated to the probability of obese individuals developing type 2 diabetes (Bento et al., 2008; Lewis et al., 2010). As a target of PI3K, P-Rex1 widely mediates insulin signalling (Balamatsias et al., 2011; Ghalali et al., 2014; Kim et al., 2011; Montero et al., 2013). P-Rex1 interacts with mTORC1, a central controller of cell growth and metabolism, through direct binding to mTOR (Hernandez-Negrete et al., 2007), and apparently can signal both up- and downstream of mTOR (Hernandez-Negrete et al., 2007; Kim et al., 2011). In 3T3-L1 adipocytes, P-Rex1 is required for the insulin-stimulated upregulation of GLUT4 from storage vesicles to the plasma membrane, promoting insulin-stimulated glucose uptake in a Rac-GEF activity dependent manner (Balamatsias et al., 2011). In human brown adipose cells, P-Rex1 regulates thermogenic potential without affecting adipocyte differentiation (Xue et al., 2015). In INS-1 832/13 pancreatic β cells, P-Rex1 is required for glucose-dependent insulin release (Thamilselvan et al., 2020). Finally, adenoviral Prex1 depletion was recently shown to protect mice from HFD-induced NAFLD (Li et al., 2022).

These findings suggested that P-Rex1 might play a role in glucose metabolism. However, it seemed unlikely that any such role would resemble P-Rex2, as P-Rex1 cannot bind and inhibit PTEN. We investigated here the role of P-Rex1 in glucose homeostasis and show that the GEF controls fasting blood glucose and insulin sensitivity through its Rac-GEF activity, but limits glucose clearance through an adaptor function involving the orphan GPCR Gpr21 in the liver.

## Results

### Prex1 limits glucose clearance, independently of its catalytic activity, and is required for the development of diet-related metabolic syndrome

To study the role of Prex1 in glucose homeostasis, we tested the fasting blood glucose levels, glucose tolerance and insulin sensitivity of *Prex1^−/−^* mice (Welch et al., 2005) throughout ageing from young adult (10 weeks) to post-reproductive ‘young-old’ age (1 year), equivalent to 20 and 60 years in humans (Dutta and Sengupta, 2016). To assess if any roles in glucose homeostasis require the catalytic Rac-GEF activity of Prex1, we generated a *Prex1^N233A/N233A^* (*Prex1^GD^*) mouse strain using CRISPR/Cas9 gene editing to introduce a N233A mutation in the catalytic DH domain that renders Prex1 catalytically inactive (GEF-dead) (Lucato et al., 2015) **(Supplemental Figure 1)**. *Prex1^GD^* mice are viable and fertile, although born at somewhat less than the expected Mendelian ratio **(Supplemental Figure 1E)**. The introduction of the N233A mutation did not affect the expression level of Prex1 protein **(Supplemental Figure 1F)**. *Prex1^GD^*mice share the white belly phenotype characteristic of *Prex1^−/−^* mice **(Supplemental Figure 1G)**, which is caused by impaired migration of melanoblasts during development (Lindsay et al., 2011), confirming that melanoblast migration depends on the Rac-GEF activity of Prex1.

*Prex1^−/−^* mice showed improved glucose tolerance compared to wild type upon glucose administration **(Figure 1A)**, whereas *Prex1^GD^* mice did not **(Figure 1B)**. Hence, Prex1 limits glucose tolerance through an adaptor function. The enhanced glucose tolerance of *Prex1^−/−^* mice persisted throughout ageing **(Figure 1C)**. Both *Prex1^−/−^* and *Prex1^GD^* mice showed lower fasting blood glucose levels, by 2-3 mM, which also persisted throughout ageing **(Figure 1A, B and Supplemental Figure 2A)**, despite having normal body weight **(Supplemental Figure 2B)**. Therefore, *Prex1^−/−^* mice have better glucose tolerance and lower fasting blood glucose than *Prex^+/+^*controls. These phenotypes are not linked mechanistically, as glucose tolerance is an adaptor function of Prex1, whereas control of fasting blood glucose depends on the Rac-GEF activity.

**Figure 1.**
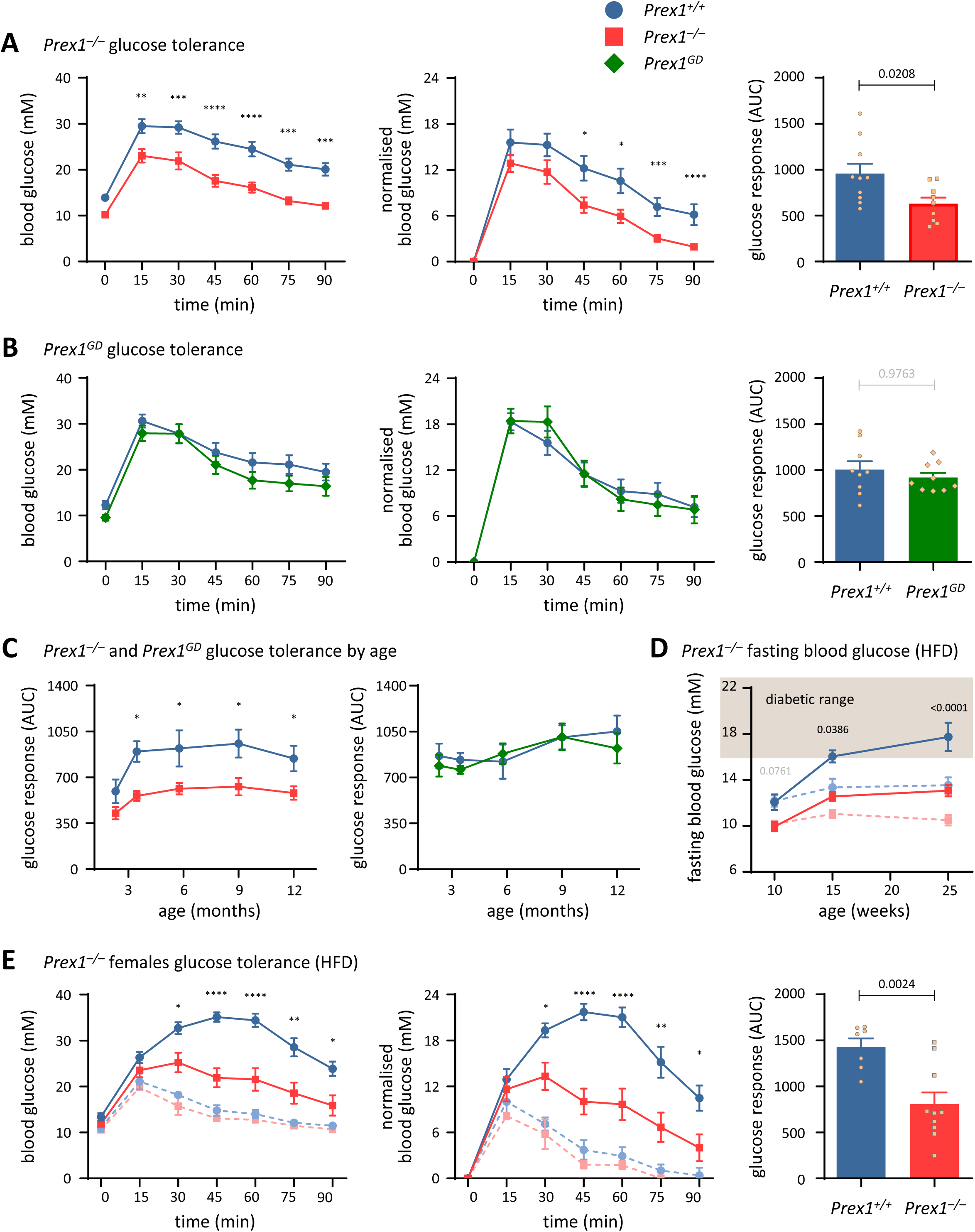
Prex1 limits glucose clearance in mice throughout ageing, independently of its GEF-activity, contributing to the diet-induced glucose intolerance and type 2 diabetes. **(A-C)** Glucose tolerance. Glucose tolerance of male (A) *Prex1^+/+^* (blue circles) and *Prex1^−/−^* (red squares) mice on chow diet, and of (B) *Prex1^+/+^* and *Prex1^GD^* (*Prex1^N233A/N233A^*, green diamonds) mice on chow diet was measured at the ages of 10 and 15 weeks, 6, 9 and 12 months. Food was withdrawn for 6 h, fasting blood glucose tested, 2 g/kg glucose injected *i.p.,* and glucose tolerance assessed at the indicated time points. (A, B) 9-month-old males. Left-hand panels: blood glucose concentration, middle: response normalised to fasting blood glucose, right: integrated normalised response (AUC). Data are mean ± SEM of mice pooled from 2 independent cohorts of 4-5 mice per genotype, 10 *Prex1^+/+^* and 10 *Prex1^−/−^* mice in (A), 9 *Prex1^+/+^* and 10 *Prex1^GD^* mice in (B); beige dots show individual mice. (C) Integrated normalised response of (A, left) and (B, right) over age. **(D)** Fasting blood glucose levels (6 h after food withdrawal) of male *Prex1^+/+^* and *Prex1^−/−^* mice on HFD from 10 weeks of age. The diabetic range is indicated. Faint symbols with stippled lines: fasting blood glucose of mice in (A) on chow diet, for reference. Data are mean ± SEM of 8 *Prex1^+/+^* and 10 *Prex1^−/−^* mice pooled from 2 independent cohorts with 4-5 mice per genotype. **(E)** Glucose tolerance of 6-months-old female *Prex1^+/+^* and *Prex1^−/−^* mice on HFD. Panels are as in (A, B). Data are mean ± SEM of 8 *Prex1^+/+^* and 10 *Prex1^−/−^*females pooled from 2 independent cohorts of 4-5 mice per genotype; beige dots show individual mice. Faint symbols with stippled lines are from one cohort of 6-months-old females on chow diet for reference. Statistics in time courses (A-E) are 2-way ANOVA with Sidak’s multiple comparisons correction; stars denote significance between genotypes for the indicated time points. Statistics in bar graphs are unpaired Student’s *t-*test; p-values in black denote significant differences, p-values in grey are non-significant.

On high-fat diet (HFD), the fasting blood glucose levels of *Prex^+/+^*control mice rose with age, reaching diabetic range in middle-age (6 months). In contrast, the fasting blood glucose of *Prex1^−/−^* mice on HFD remained below the diabetic range **(Figure 1D)**, despite gaining diet-related excess body weight and body fat to the same extent as *Prex^+/+^* controls **(Supplemental Figure 2C-D)**. Similarly, *Prex1^−/−^*mice were protected from developing HFD-related glucose intolerance **(Figure 1E)**. Hence Prex1 deficiency protects mice from developing diet-dependent metabolic syndrome and type 2 diabetes. The increased glucose tolerance and reduced fasting blood glucose were seen both in males and females, with some diet-dependent differences, even though females generally displayed better glucose tolerance than males **(Figure 1A,E**, and data not shown).

### Prex1 is required for insulin sensitivity, through its Rac-GEF activity

Administration of insulin induced a drop in blood glucose levels in both *Prex^+/+^* and *Prex1^−/−^* mice. However, blood glucose did not decrease to the same extent in *Prex1^−/−^* mice, and *Prex1^−/−^* mice could not sustain their response to insulin, returning to baseline blood glucose faster **(Figure 2A)**. Hence, Prex1 is required for insulin sensitivity, particularly for maintaining the response to insulin. This phenotype was age-dependent, as insulin resistance was seen in 10-month-old but not 5-month-old *Prex1^−/−^* mice **(Figure 2A** and data not shown). *Prex1^GD^* mice were more severely insulin resistant than *Prex1^−/−^* mice, their blood glucose hardly decreasing in response to insulin, and their phenotype was already apparent at the age of 5 months **(Figure 2A and Supplemental Figure 3A)**. Hence, Prex1 is required for insulin sensitivity, through its Rac-GEF activity. The finding that *Prex1^GD^* mice had more severe insulin resistance than *Prex1^−/−^* mice suggests that catalytically inactive Prex1 may have a dominant-negative effect on insulin responses.

**Figure 2.**
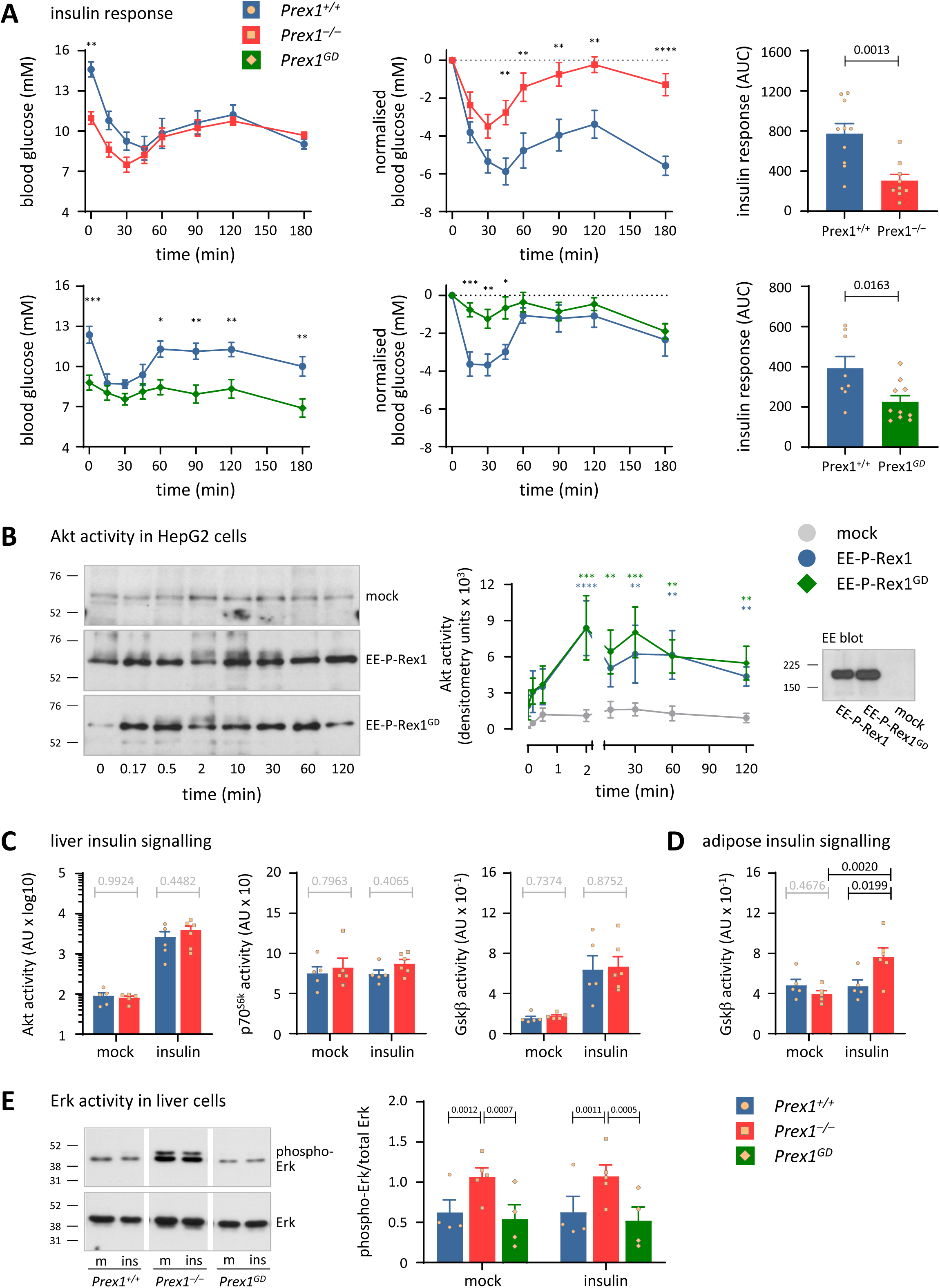
Prex1 is required for insulin sensitivity through its Rac-GEF activity, and limits Erk activity in the liver independently of its catalytic activity. **(A)** Insulin sensitivity of *Prex1^+/+^* (blue circles), *Prex1^−/−^*(red squares, top), and *Prex1^GD^* (green diamonds, bottom) mice. Mice were fasted for 4 h, fasting blood glucose was measured, 0.75 IU/kg insulin (top) or 0.375 IU/kg insulin (bottom) *s.c.* injected, and blood glucose tested at the indicated timepoints. Left-hand panel: blood glucose concentration, middle: response normalised to fasting blood glucose, right: integrated normalised response (AAC; beige dots show individual mice). The *Prex1^−/−^* mice in Figure 1A were tested at the age of 10 months (43 weeks). Data are mean ± SEM of 10 *Prex1^+/+^*and 10 *Prex1^−/−^* mice pooled from 2 independent cohorts of 4-5 mice per group. The *Prex1^GD^* mice in Figure 1B were tested at the age of 5 months (21 weeks). Data are mean ± SEM of 9 *Prex1^+/+^*and 10 *Prex1^GD^* mice pooled from 2 independent cohorts of 4-5 mice per group. Statistics in time courses (A-C) are 2-way ANOVA with Sidak’s multiple comparisons correction; stars denote significance between genotypes for the indicated time points. Statistics in bar graphs are unpaired Student’s *t-*test. **(B)** HepG2 cells expressing wild type P-Rex1 (blue circles) or GEF-dead P-Rex1 (green diamonds), or mock-transfected (grey circles), were serum-starved, stimulated with 25 nM insulin for the indicated periods of time, and total cell lysates western blotted for phospho-Akt (S473) and quantified by densitometry. Blots shown are from one representative experiment. EE blot shows expression of wild type and GEF-dead P-Rex1. Data are mean ± SEM of 6 independent experiments, normalized to the mock-transfected 0-time point. Statistics are 2-way ANOVA on square-root transformed raw data with Sidak’s multiple comparisons correction. Blue stars denote differences between mock and wild type P-Rex1, green stars between mock and GEF-dead P-Rex1. **(C,D)** Mesoscale analysis of insulin signalling in mouse liver (C) and white adipose tissue (D). 10-week-old *Prex1^+/+^* (blue bars, circles) and *Prex1^−/−^* (red bars, squares) mice were fasted for 4 h, injected *s.c.* with 0.75 IU/kg insulin, or mock-treated, culled humanely after 15 min, tissues retrieved, homogenised, and active and total levels of Akt, p70^S6K^, and GSK3β detected by Mesoscale analysis. Data are mean ± SEM of 5-6 mice per group; beige symbols show individual mice. Statistics are 2-way ANOVA with Sidak’s multiple comparisons correction on log-transformed raw data; p-values in black are significant; p-values in grey are non-significant. **(E)** Erk activity. 1 × 10^6^ liver cells isolated from 15-week-old *Prex1^+/+^* (blue bars, circles), *Prex1^−/−^* (red bars, squares), and *Prex1^GD^* (green bars, diamonds) mice were stimulated with 100 nM insulin for 10 min at 37°C, and lysates analysed by western blotting with phospho-Erk and total Erk antibodies. Blots were quantified using Fiji. Data are mean ± SEM of 4-5 experiments and are expressed as phospho-Erk/total Erk signal; beige symbols show individual experiments. Statistics are 2-way ANOVA with Sidak’s multiple comparisons correction.

In summary, Prex1 limits glucose tolerance through an adaptor function and controls fasting blood glucose levels and insulin sensitivity through its catalytic Rac-GEF activity.

### Prex1 is not required for insulin signalling throug*h Akt, p70^S6k^ and Gskβ* in metabolic tissues but limits Erk activity in primary liver cells

Prex1 mediates insulin signalling in a range of cell types (Balamatsias et al., 2011; Ghalali et al., 2014; Kim et al., 2011; Montero et al., 2013). As Prex1 was required for insulin sensitivity *in vivo*, we asked if it controls insulin signalling in cells and tissues relevant to glucose homeostasis. First, we overexpressed human wild type or GEF-dead EE-tagged P-Rex1 in the human hepatocyte cell line HepG2, stimulated the cells with insulin, and assessed Akt (phospho-S473) activity. Expression of both wild type and GEF-dead P-Rex1 increased insulin-stimulated Akt activity **(Figure 2B)**. Hence, Prex1 can promote insulin signalling when overexpressed, independently of its Rac-GEF activity.

To assess insulin signalling in primary mouse tissues, we challenged *Prex^+/+^* and *Prex1^−/−^*mice with insulin, retrieved liver, skeletal muscle, and white adipose tissues after 15 min, and analysed the activities of Akt, p70^S6k^ and Gskβ in tissue lysates by Mesoscale analysis. Insulin challenge increased the activities of Akt and Gskβ in all tissues, and raised p70^S6k^ activity in skeletal muscle and adipose tissue but not the liver **(Figure 2C and Supplemental Figure 3B,C)**. However, all activities were identical between *Prex^+/+^* and *Prex1^−/−^*, except for somewhat elevated Gskβ activity in insulin-stimulated *Prex1^−/−^* adipose tissue **(Figure 2C,D and Supplemental Figure 3B,C)**. Hence, Prex1 is not required for insulin-stimulated Akt and p70^S6k^ signalling in metabolic tissues, but limits insulin-stimulated Gskβ activity in adipose tissue.

We also assessed Erk activity in primary liver cells isolated from *Prex^+/+^*, *Prex1^−/−^* and *Prex1^GD^* mice. *Prex1^−/−^* liver cells showed constitutively increased Erk activity compared to *Prex^+/+^* and *Prex1^GD^* cells, and this Erk activity was not affected by insulin stimulation under the conditions tested **(Figure 2E)**. Hence, Prex1 constitutively limits Erk activity in liver cells, independently of its catalytic Rac-GEF activity.

In summary, Prex1 can promote insulin-stimulated Akt signalling in HepG2 cells but is not required for Akt or p70^S6k^ signalling in primary metabolic tissues. However, Prex1 limits constitutive Erk activity in liver cells, independently of its catalytic activity, and it limits insulin-stimulated Gskβ activity in adipose tissue. This signalling pathway analysis does not obviously explain the GEF-activity dependent roles of Prex1 in fasting blood glucose and insulin sensitivity, but its GEF-activity independent function in limiting Erk activity might contribute to its GEF-activity independent role in glucose clearance.

### Prex1 does not control glucose excretion, plasma insulin, or liver glycogen

To begin to investigate possible causes of the lower fasting blood glucose level and improved glucose tolerance, we tested *Prex1^−/−^* mice in metabolic cages. In line with their normal body weight and body fat, *Prex1^−/−^* mice consumed normal amounts of food and water and produced normal amounts urine and faeces, regardless of age, sex, and diet **(Supplemental Figure 4)**. We asked whether *Prex1^−/−^* mice clear glucose more efficiently by excreting it more. However, the amount of glucose excreted in urine upon glucose administration was normal **(Figure 3A)**. This normal level of excretion implied that the increased glucose tolerance of *Prex1^−/−^*mice is caused by more efficient glucose clearance from the blood stream into tissues. To test if *Prex1^−/−^* mice clear glucose better because of increased insulin levels, we measured plasma insulin. However, plasma insulin was normal both in fasted *Prex1^−/−^* mice and upon acute glucose challenge, under conditions where glucose was cleared faster from the blood stream, although a slight raise in plasma insulin was observed at late timepoints, after the glucose was largely cleared **(Figure 3B)**. Similarly, plasma glucagon and adiponectin levels were normal both in fasting mice and upon glucose administration **(Supplemental Figure 5A-B)**. In line with the normal plasma insulin, glucose-stimulated secretion of insulin by pancreatic islets isolated from *Prex1^−/−^* mice was also normal **(Supplemental Figure 5C)**. The only difference in hormone levels we observed was a raise in plasma glucagon in *Prex1^−/−^* mice upon insulin administration **(Supplemental Figure 5D)**. The latter may in part explain the inability of *Prex1^−/−^* mice to sustain low blood glucose levels after insulin challenge.

**Figure 3.**
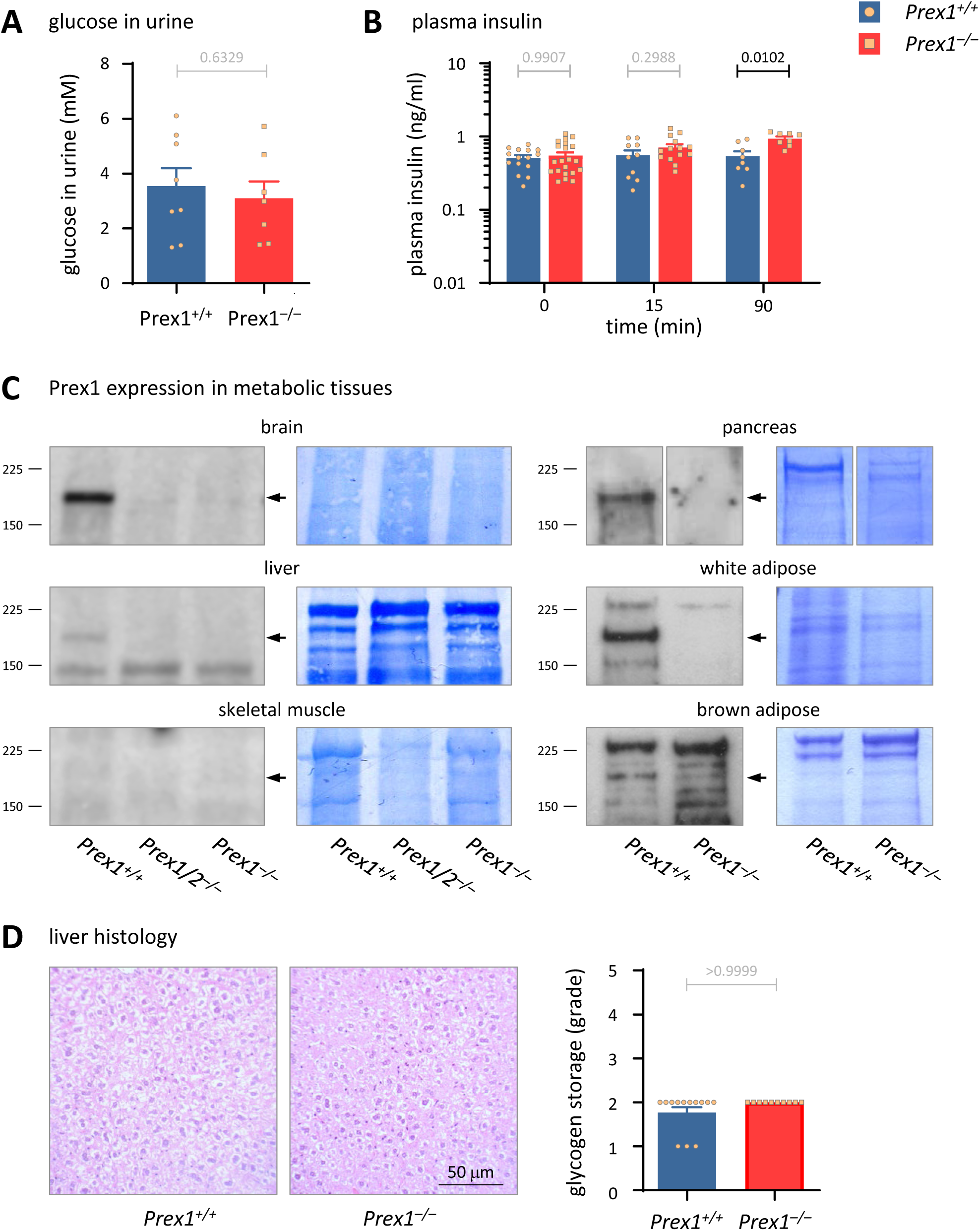
Prex1 is expressed in metabolic tissues, without controlling glucose excretion, plasma insulin, or liver glycogen. **(A)** Glucose excretion. 15-week-old *Prex1^+/+^* (blue bars, circles) and *Prex1^−/−^* (red bars, squares) males on chow diet were fasted for 6 h before 2 g/kg glucose was *i.p.* injected. Urine was collected 90 min later and glucose concentration assessed by ELISA. Data are mean ± SEM of 8 Prex1^+/+^ and 7 Prex1^−/−^ mice pooled from 2 independent cohorts of 3-4 mice per genotype; beige symbols show individual mice. Statistics are unpaired Student’s *t-*test; p-values in grey are non-significant. **(B)** Plasma insulin. Insulin in blood plasma of 15-week-old *Prex1^+/+^* and *Prex1^−/−^*males on chow diet fasted for 6 h (0 time), or fasted and then challenged with *i.p.* 2 g/kg glucose for 15 or 90 min, was analysed by ELISA. Data are mean ± SEM of mice pooled from 3 independent cohorts (4 for 0 time); beige symbols show individual mice. Statistics are 2-way ANOVA on square root-transformed raw data with Sidak’s multiple comparisons correction; p-values in black denote significant differences, p-values in grey are non-significant. **(C)** Prex1 expression in metabolic tissues. Total lysates of the indicated metabolic tissues from *Prex1^+/+^*, *Prex1^−/−^*, and *Prex1^−/−^Prex2^−/−^* mice were western blotted with P-Rex1 antibody. Blots are representative of 3 independent experiments. Coomassie staining was done to control for protein loading. **(D)** Liver histology. Livers from 10-week-old *Prex1^+/+^*(blue bars, circles) and *Prex1^−/−^* (red bars, squares) males on chow diet were stained with H&E, and glycogen storage was assessed by a pathologist. Data are mean ± SEM of mice pooled from 2 independent cohorts; beige symbols show individual mice. Statistics are chi-square test.

To search for tissues into which *Prex1^−/−^* mice might preferentially clear blood glucose, we first western blotted mouse tissues for Prex1 expression. Prex1 was expressed in brain, liver, white and brown adipose tissue, but was barely detectable in skeletal muscle **(Figure 3C)**. As previously described, the livers of *Prex1^−/−^* mice were smaller than controls (Welch et al., 2005), regardless of age and sex, and the same phenotype was seen in *Prex1^GD^* mice **(Supplemental Figure 6A)**. All other organs were of normal size (data not shown). To test if the increased glucose clearance in *Prex1^−/−^* mice may stem from preferential glycogen storage in the liver, we performed histological analysis. However, liver glycogen was normal **(Figure 3D)**. Histological analysis of skeletal muscle and white and brown adipose tissues also revealed no abnormalities (data not shown). Expression levels of the liver enzymes Pepck and glucose 6-phosphatase were normal, both in fasted animals and upon insulin administration **(Supplemental Figure 6B)**.

### Prex1 limits glucose uptake, Glut2 surface levels and ATP production in liver cells, independently of its GEF-activity

As our initial tissue analysis gave no clues as to where blood glucose may be cleared, with the possible exception of elevated Erk activity in liver cells, we determined glucose uptake empirically. We isolated cells from the liver, skeletal muscle, and various adipose tissues of *Prex^+/+^*, *Prex1^−/−^* and *Prex1^GD^* mice and measured the constitutive and insulin-stimulated uptake of tritiated 2-deoxyglucose (^3^H-2-DOG). *Prex1^−/−^* liver cells showed constitutively increased glucose uptake, whereas glucose uptake was normal in *Prex1^GD^* liver cells **(Figure 4A)**. A small, constitutive increase in glucose uptake was also seen in *Prex1^−/−^* skeletal muscle cells **(Supplemental Figure 7A)**, whereas glucose uptake into mature *Prex1^−/−^* visceral white adipocytes, subcutaneous white adipocytes, and interscapular brown adipocytes was normal **(Supplemental Figure 7B-D)**. Insulin stimulation did not raise glucose uptake in liver cells **(Figure 4A)** but did in *Prex^+/+^* skeletal muscle cells, up to the level seen constitutively in *Prex1^−/−^* **(Supplemental Figure 7A)**. Insulin stimulation also increased glucose uptake into adipose cells, without revealing any defects in *Prex1^−/−^* **(Supplemental Figure 7B-D)**. Hence, Prex1 limits glucose uptake into liver cells through an adaptor function, and to some degree in skeletal muscle cells, but does not affect glucose uptake into adipocytes. The substantial increase in glucose uptake into liver cells can explain the increased glucose clearance seen in *Prex1^−/−^* mice *in vivo*.

**Figure 4.**
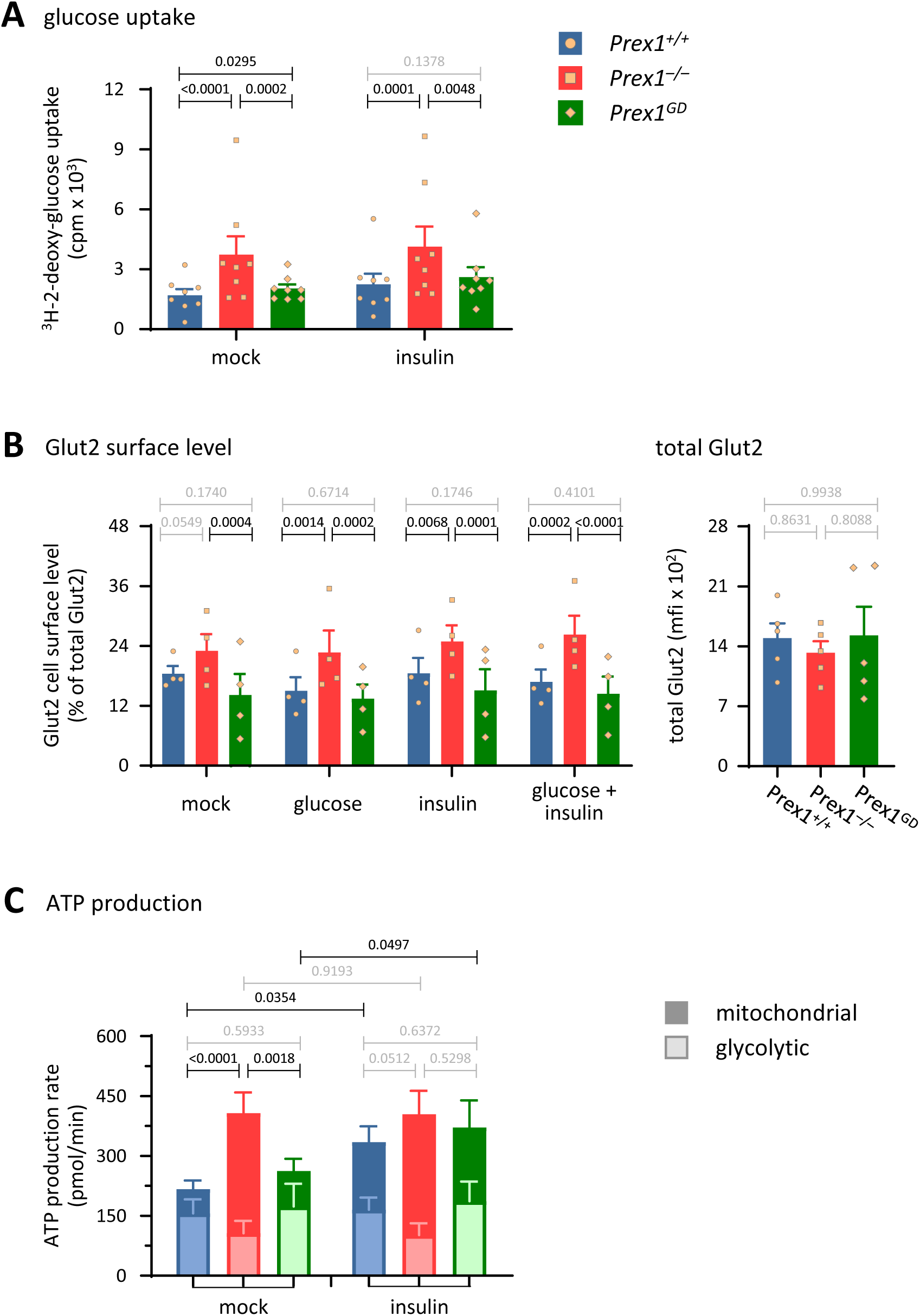
Prex1 limits glucose uptake, Glut2 surface levels and ATP production in liver cells, independently of its GEF-activity. **(A)** Glucose uptake. Liver cells from 15 week-old *Prex1^+/+^* (blue bars, circles), *Prex1^−/−^* (red bars, squares) and *Prex1^GD^* (green bars, diamonds) males were stimulated with 100 nM insulin for 10 min at 37°C, or were mock-stimulated, followed by the addition of 50 μM 2-DOG, 0.25 μCi ^3^H-2-DOG for a further 60 min. Cells were washed, lysed, and glucose uptake was measured by scintillation counting. Data are mean ± SEM of 8 independent experiments; beige symbols show individual experiments. Statistics are 2-way ANOVA on log-transformed raw data with Sidak’s multiple comparisons correction; p-values in black denote significant differences, p-values in grey are non-significant. **(B)** Glut2 cell surface level. Left: Liver cells from mice as in (A) were stimulated with 5 mM glucose or 100 nM insulin for 10 min at 37°C, or were mock-stimulated, or stimulated with 100 nM insulin for 10 min and 5 mM glucose for another 30 min, stained with Glut2 antibody, analysed by flow cytometry, and the mean fluorescence intensity (mfi) of Glut2 surface level expressed as % of total Glut2. Data are mean ± SEM of 4 independent experiments; beige symbols show individual experiments. Statistics are 2-way ANOVA with Sidak’s multiple comparisons correction. Right: Total Glut2 was measured in the same way except in permeabilised cells. Data are mean ± SEM of 5 independent experiments; beige symbols show individual experiments. Statistics are one-way ANOVA with Tukey’s multiple comparisons correction. **(C)** ATP production. Liver cells from mice as in (A) were analysed by Seahorse assay in the presence or absence of 100 nM insulin to quantify ATP production from glycolysis (light bars) and mitochondrial respiration (dark bars). Data are mean ± SEM of 5-8 independent experiments. Statistics are 2-way ANOVA with Sidak’s multiple comparisons correction for total ATP production.

Glucose enters liver cells mainly through glucose transporter Glut2, which is sensitive to glucose levels but not insulin (Navale and Paranjape, 2016; Thorens, 2015). We measured the total and cell surface levels of Glut2 by flow cytometry. The cell surface level of Glut2 was constitutively raised in *Prex1^−/−^*liver cells, regardless of stimulation with glucose or insulin **(Figure 4B)**. In contrast, the cell surface level of Glut2 was normal in *Prex1^GD^* liver cells under all conditions tested, and the total cellular level of Glut2 was normal in all genotypes **(Figure 4B)**. In contrast to Glut2, the surface level of Glut1, another insulin-independent glucose transporter in liver cells, was normal **(Supplemental Figure 8)**. These data suggested that *Prex1^−/−^* liver cells may take up glucose more readily because of the increased cell surface localisation of Glut2. As total Glut2 levels were normal, the data suggested furthermore that Prex1 might regulate Glut2 trafficking, through an adaptor function.

As glucose entered *Prex1^−/−^* liver cells more readily, but liver glycogen storage appeared normal, we reasoned that the excess glucose was likely metabolised to ATP. We tested ATP production by Seahorse assay, distinguishing between glycolytic and mitochondrial ATP. Indeed, *Prex1^−/−^* liver cells showed constitutively increased mitochondrial ATP production, whereas mitochondrial ATP production was normal in *Prex1^GD^* liver cells, and glycolytic ATP production was normal in both genotypes **(Figure 4C)**. Insulin stimulation increased mitochondrial ATP production in *Prex^+/+^* and *Prex1^GD^* cells, to the level seen constitutively in *Prex1^−/−^* cells, but did not affect ATP production further in *Prex1^−/−^* cells **(Figure 4C)**.

Hence, *Prex1^−/−^*mice have increased glucose clearance into liver cells, likely *via* the raised cell surface level of Glut2, and this results in increased glucose metabolism through mitochondrial ATP production, all independently of the catalytic Rac-GEF activity of Prex1.

### Prex1 controls glucose uptake and mitochondrial ATP production in liver cells through Gpr21

We recently showed that P-Rex1 has one other adaptor function, in addition to the control of glucose clearance, which is to limit the agonist-induced trafficking of G protein coupled receptors (GPCRs) (Baker, Hampson et al., manuscript under review at Cell Reports). We hypothesised that both processes might be linked, and Prex1 may control glucose uptake and metabolism in liver cells through its adaptor function in GPCR trafficking.

One GPCR implicated in glucose uptake into liver cells is Gpr21, a constitutively active Gα_q_-coupled orphan GPCR. Initial reports suggesting that Gpr21 deficiency improves the glucose tolerance and insulin sensitivity of mice on HFD (Gardner et al., 2012; Osborn et al., 2012) were drawn into question when it was shown that Rapgap1 was accidentally co-deleted (Wang et al., 2016). However, a recent, independently generated *Gpr21^−/−^* mouse with normal Rapgap1 expression also shows improved glucose tolerance (Riddy et al., 2021). Overexpression of GPR21 in Hek293 cells stimulates phospholipase C and Mapk pathway activities but inhibits insulin signalling and glucose uptake (Leonard et al., 2016), whereas knockdown of GPR21 in HepG2 cells inhibits phospholipase C but promotes insulin signalling and glucose uptake (Kinsella et al., 2021). Hence, Gpr21 limits glucose uptake in liver cells. Few tools are available for studying Gpr21, with the exception of western blotting antibodies and a small-molecule inverse agonist, GRA2, which blocks the constitutive activity of Gpr21, and therefore the inhibitory action of the GPCR on insulin signalling and glucose uptake (Kinsella et al., 2021; Leonard et al., 2016). We verified the specificity of GRA2 by siRNA-mediated knockdown of GPR21 in HepG2 cells **(Supplemental Figure 9A-C)**. In HepG2 cells treated with non-targeting control siRNA, GRA2 increased glucose uptake. Treatment of HepG2 cells with GPR21 siRNA also enhanced glucose uptake, but GRA2 had no effect in these GPR21 siRNA-treated cells **(Supplemental Figure 9C)**, as expected (Kinsella et al., 2021).

Prex1 deficiency did not affect Gpr21 expression in mouse liver cells **(Supplemental Figure 9D)**. To test if Prex1 controls the trafficking of Gpr21, as it does with other GPCRs, we fractionated *Prex^+/+^* and *Prex1^−/−^* liver cells by differential centrifugation under detergent-free conditions and isolated endosomes from the 10000 × g supernatant by Eea1 immunoprecipitation (Gosney, 2016), prior to ultracentrifugation to obtain the plasma membrane fraction. Use of Eea1 as a marker for early endosomes and Kras as a marker for the plasma membrane showed that the two fractions were pure. Western blotting revealed increased endosomal localisation of Gpr21 in *Prex1^−/−^* liver cells **(Figure 5A)**. As total cellular levels of Gpr21 were normal, we can conclude that Prex1 controls the trafficking of Gpr21 by limiting the endosomal localisation of the GPCR. We could not detect a band of the expected size for Gpr21 in the plasma membrane fraction, but a band of slightly higher molecular weight was taken to be Gpr21 and was evenly localised the between *Prex^+/+^* and *Prex1^−/−^***(Figure 5A)**.

**Figure 5.**
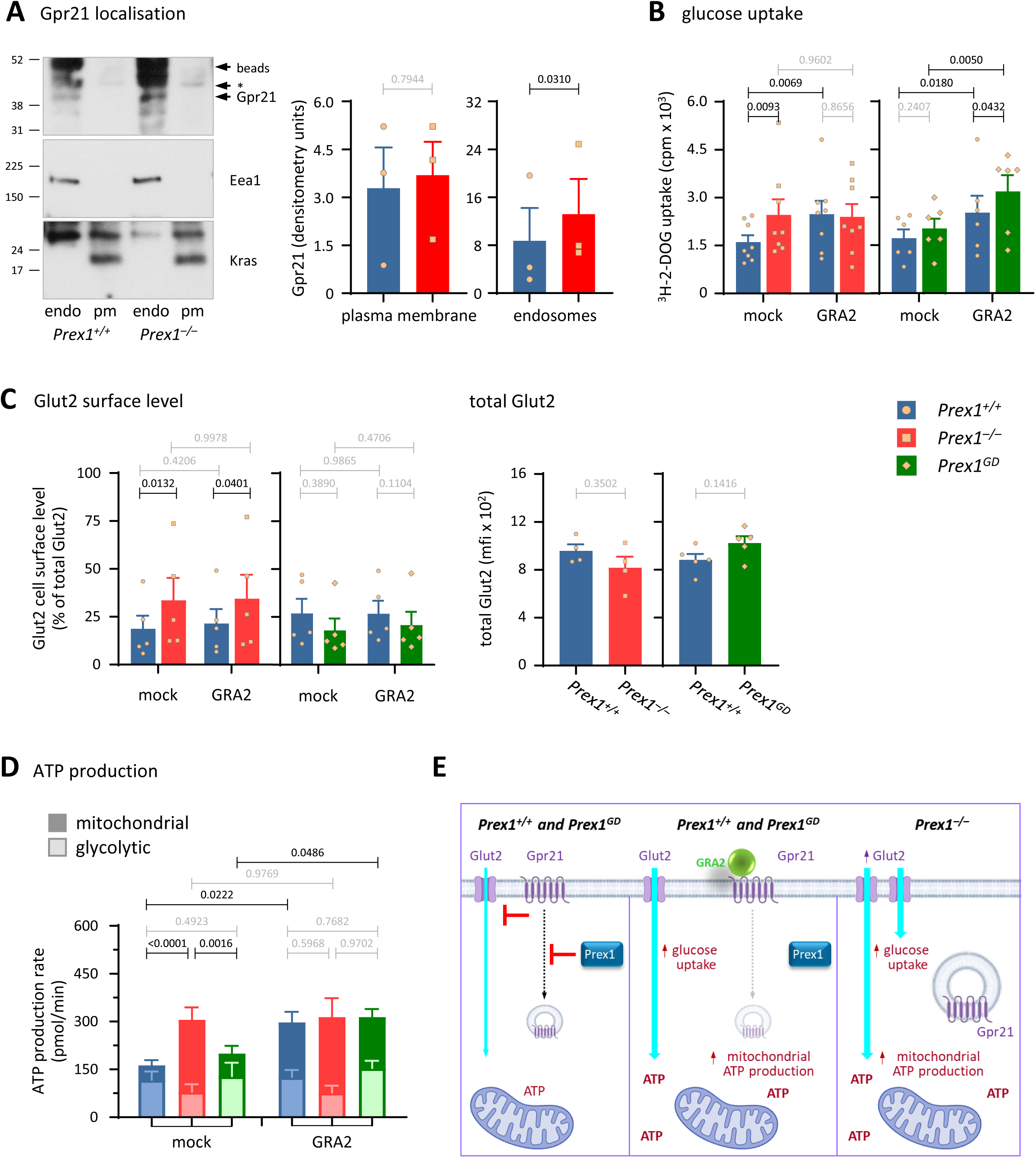
Prex1 limits glucose uptake and ATP production in liver cells through Gpr21, independently of its GEF activity. **(A)** Gpr21 localisation. Liver cells from 16-week-old *Prex1^+/+^*(blue bars, circles) and *Prex1^−/−^* (red bars, squares) males were fractionated by differential centrifugation under detergent-free conditions. Endosomes (endo) were isolated from the 10000 × g supernatant by Eea1 immunoprecipitation, prior to ultracentrifugation to obtain the plasma membrane fraction (pm). Fractions were western blotted for Gpr21. Eea1 was used as a marker for early endosomes and Kras as a marker for the plasma membrane. The same cell equivalents of endosomal and plasma membrane fractions were loaded. The asterisk denotes a band of higher than expected MW in the plasma membrane fraction taken to be Gpr21. Western blots shown are from 1 experiment representative of 3. Blots were quantified by Fiji densitometry. Data are mean ± SEM of 3 independent experiments. Statistics are 2-way ANOVA with Sidak’s multiple comparisons correction. **(B)** Glucose uptake. Liver cells from 15-week-old *Prex1^+/+^*, *Prex1^−/−^* and *Prex1^GD^* (green bars, diamonds) males were treated with 30 μM GRA2, an inverse agonist of Gpr21, for 3 h at 37°C, or were mock-stimulated, followed by the addition of 50 μM 2-DOG, 0.25 μCi ^3^H-2-DOG for a further 60 min. Cells were washed, lysed, and glucose uptake was measured by scintillation counting. Data are mean ± SEM of 9 independent experiments for *Prex1^−/−^* and 6 for *Prex1^GD^*; beige symbols show individual experiments. Statistics are 2-way ANOVA on square root-transformed raw data with Sidak’s multiple comparisons correction; p-values in black denote significant differences, p-values in grey are non-significant. **(C)** Glut2 cell surface level. Left: Liver cells from mice as in (A) were treated with 30 μM GRA2 for 3 h at 37°C, or mock-stimulated, stained with Glut2 antibody, and analysed by flow cytometry. The mfi of Glut2 at the cell surface was expressed as % of total Glut2. Data are mean ± SEM of 5 independent experiments for *Prex1^−/−^* and 5 for *Prex1^GD^*; beige symbols show individual experiments. Statistics are 2-way ANOVA on square-root transformed raw data with Sidak’s multiple comparisons correction. Right: The total cellular level of Glut2 was measured in the same manner except with permeabilised cells. Data are mean ± SEM of 4 independent experiments for *Prex1^−/−^* and 5 for *Prex1^GD^*; beige symbols show individual experiments. Statistics are paired t-test; p-values in grey are non-significant. **(D)** ATP production. Liver cells from mice as in (A) were analysed by Seahorse assay in the presence or absence of 30 μM GRA2 to quantify ATP production from glycolysis (light bars) and mitochondrial respiration (dark bars). Data are mean ± SEM of 5-8 independent experiments. Statistics are 2-way ANOVA with Sidak’s multiple comparisons correction for total ATP production. **(E)** Model. Schematic generated in part using BioRender.

To test if Prex1 regulates glucose uptake through Gpr21, we treated liver cells isolated from *Prex^+/+^*, *Prex1^−/−^*and *Prex1^GD^* mice with GRA2 and measured the uptake of ^3^H-2-DOG. As seen before, glucose uptake was constitutively increased in *Prex1^−/−^*liver cells, whereas it was normal in *Prex1^GD^* liver cells. GRA2 treatment increased glucose uptake in *Prex^+/+^* and *Prex1^GD^* liver cells, but not in *Prex1^−/−^* cells **(Figure 5B)**. Hence, Prex1 limits glucose uptake into liver cells through an adaptor function that involves Gpr21.

Knockdown of GPR21, or GRA2 treatment, increases the cell surface localisation of GLUT2 in HepG2 cells (Kinsella et al., 2021). We did not see this in primary liver cells from *Prex^+/+^*, *Prex1^−/−^* or *Prex1^GD^* mice **(Figure 5C)**. *Prex1^−/−^* liver cells showed the expected constitutively raised cell surface Glut2, but GRA2 had no effect in any genotype. Prex1 limits the internalisation of active GPCRs (Baker, Hampson et al., manuscript under review at Cell Reports), and as Gpr21 is constitutively active, it seems likely that Prex1 controls Gpr21 by preventing its internalisation into endosomes.

To investigate links between Prex1 and Gpr21 in ATP production, we tested the effect of GRA2 on glycolytic and mitochondrial ATP production in liver cells from *Prex^+/+^*, *Prex1^−/−^* or *Prex1^GD^*mice by Seahorse assay. *Prex1^−/−^* liver cells showed the expected constitutively increased mitochondrial ATP production. However, GRA2 treatment increased mitochondrial ATP production in *Prex^+/+^* and *Prex1^GD^* cells, but not in *Prex1^−/−^* cells **(Figure 5D)**. Therefore, Prex1 controls glucose uptake and glucose metabolism in liver cells through Gpr21.

We propose a mechanism whereby Prex1 limits the internalisation of constitutively-active Gpr21 through its adaptor function in GPCR trafficking, thus allowing Gpr21 to inhibit glucose uptake. When Gpr21 activity is blocked by GRA2, the inhibitory effect of the receptor on glucose uptake is removed, and Prex1 does not affect the trafficking of Gpr21 in this inactive state. In *Prex1^−/−^* liver cells, the constitutively active Gpr21 is internalised as the Prex1-dependent block of receptor endocytosis is removed, and thus the Gpr21-dependent block of glucose uptake is also removed, and GRA2 cannot affect the internalised receptor **(Figure 5E)**.

### Prex1 limits mitochondrial membrane potential in liver cells

The Seahorse assays showed that Prex1 limits mitochondrial but not glycolytic ATP production. To investigate if Prex1 affects ATP production by influencing mitochondrial membrane potential, we stained liver cells from *Prex^+/+^*, *Prex1^−/−^* and *Prex1^GD^* mice with MitoTracker Green (MTG) to quantify mitochondria and with tetramethyl-rhodamine ethyl ester (TMRE), which reports mitochondrial membrane potential, and assessed the cells by flow cytometry and by widefield fluorescence live-imaging **(Figure 6)**.

**Figure 6:**
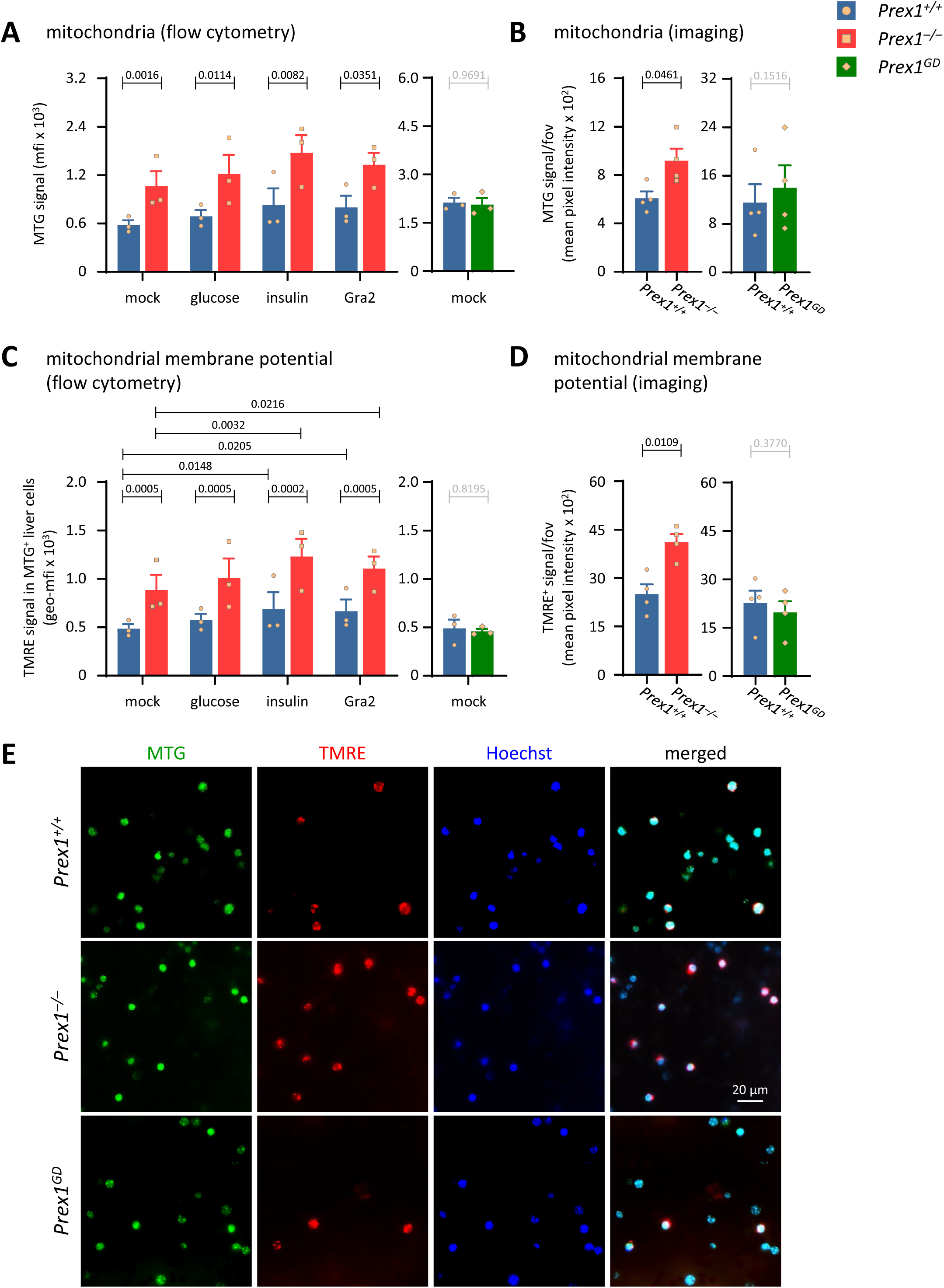
Prex1 limits mitochondrial membrane potential in liver cells. **(A-E)** Mitochondrial membrane potential. Liver cells from 15-week-old *Prex1^+/+^* (blue bars, circles), *Prex1^−/−^* (red bars, squares) or *Prex1^GD^*(green bars, diamonds) males were stimulated with 5 mM glucose or 100 nM insulin, or were treated with 30 μM GRA2 for 30 min at 37°C, or mock-stimulated, as indicated. (A, C) Flow cytometry. Cells were stained with cell-permeable MitoTracker Green (MTG) and TMRE and analysed by flow cytometry for MTG signal (A) and for TMRE signal within MTG^+^ cells (C). Data are mean ± SEM of 3 independent experiments per genotype; beige symbols show individual experiments. Statistics are 2-way ANOVA on square-root transformed raw data with Sidak’s multiple comparisons correction. (B, D, E) Imaging. Mock-stimulated liver cells were stained with MTG, TMRE, and Hoechst 33342, imaged by widefield fluorescence microscopy, and analysed for MTG signal (B) and TMRE signal (D) per field of view (fov) by Fiji. Representative images are shown in (E). Data are mean ± SEM of 4 independent experiments per genotype; beige symbols show individual experiments. Statistics are paired t-test.

*Prex1^−/−^* liver cells showed increased MTG signal under all conditions tested, both by flow cytometry and by imaging **(Figure 6 A,B,E**, and data not shown). In contrast, MTG signal was normal in *Prex1^GD^*liver cells under all conditions tested. Therefore, Prex1 limits the number or size of mitochondria in liver cells, independently of its catalytic Rac-GEF activity.

*Prex1^−/−^* liver cells also showed increased TMRE signal under all conditions tested, which demonstrates that Prex1 limits mitochondrial membrane potential. In contrast, mitochondrial membrane potential was normal in *Prex1^GD^* liver cells under all conditions tested **(Figure 6C-E**, and data not shown). Hence, Prex1 limits mitochondrial membrane potential in liver cells, independently of its catalytic activity. Stimulation with insulin, or treatment with GRA2, increased TMRE signal both in *Prex^+/+^* and *Prex1^−/−^* mice **(Figure 6C)**, and data not shown), suggesting that Prex1 deficiency does not saturate the mitochondrial membrane potential, and that insulin and Gpr21 also regulate mitochondrial activity at least in part independently of Prex1.

### Prex1 controls mitochondrial morphology in liver cells

To assess if the altered mitochondrial activity in *Prex1^−/−^*liver cells is reflected by morphological changes, we analysed the livers of *Prex^+/+^*, *Prex1^−/−^* and *Prex1^GD^*mice by focussed ion beam scanning electron microscopy (FIB-SEM). *Prex1^−/−^* liver cells had fewer but larger mitochondria than *Prex^+/+^* and *Prex1^GD^*, including mega-mitochondria **(Figure 7A,B and Supplemental Figure 10)**. Therefore, Prex1 controls mitochondrial morphology and limits mitochondrial membrane potential in liver cells, independently of its Rac-GEF activity.

**Figure 7.**
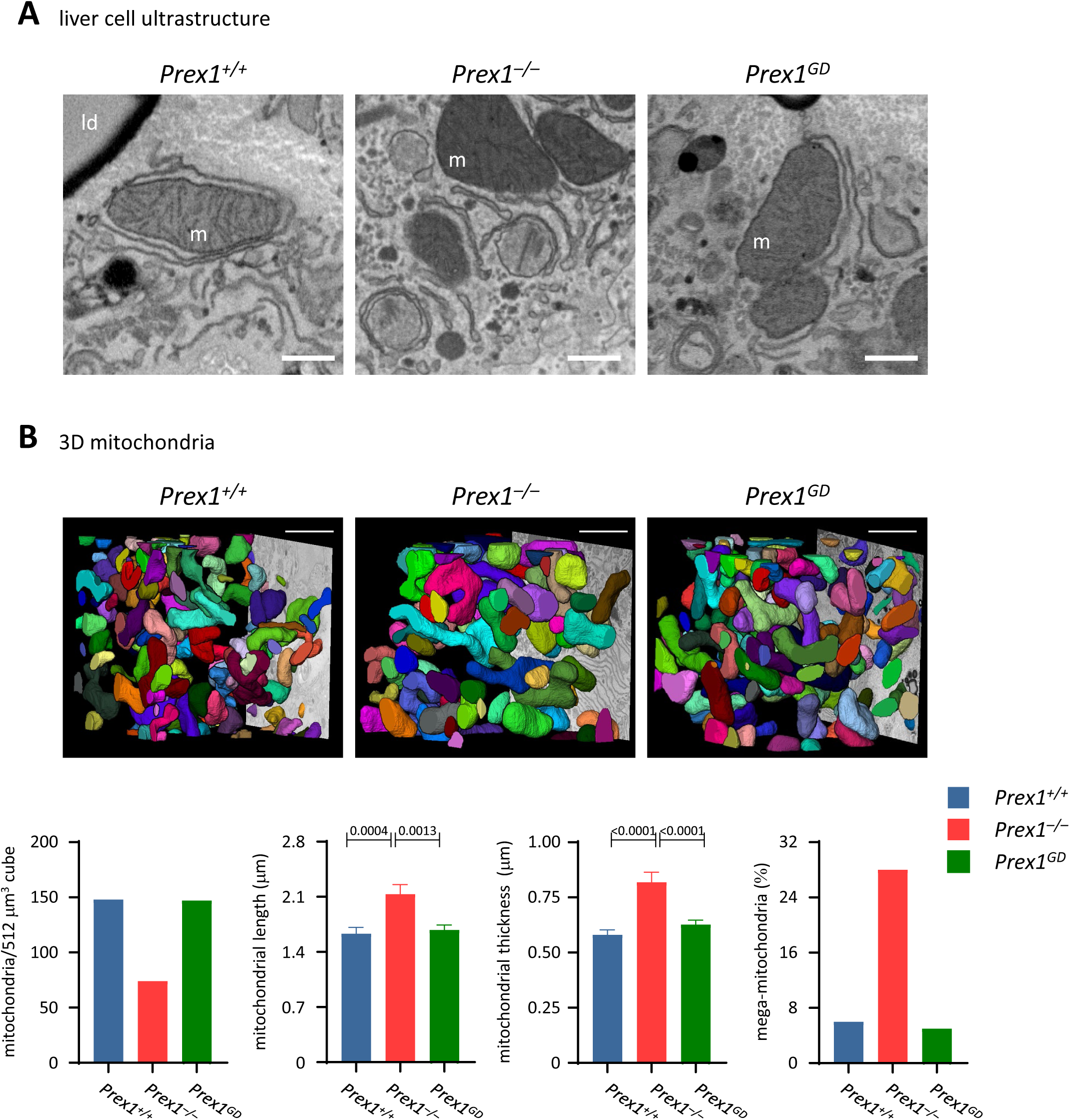
Prex1 controls mitochondrial morphology in liver cells. FIB-SEM ultrastructure of liver from 15-week-old male *Prex1^+/+^*, *Prex1^−/−^*, and *Prex1^GD^*mice. **(A)** Representative XY slices of volumetric EM data sets. Mitochondria (m) and lipid droplets (ld) are indicated. Representative images are part of a larger set shown in Supplemental Figure 10. Scale bars are 0.5 µm. **(B)** Volumetric images of mitochondria in a cube of 512 μm^3^ (8 × 8 × 8 μm). Data are the mean number of mitochondria within the cube (left; 148 *Prex1^+/+^*, 74 *Prex1^−/−^*, and 147 *Prex1^GD^*), their mean length and thickness (mean ± SEM, middle panels), and % mega-mitochondria (>1 μm length and >1 μm width) among them (right). Statistics are one-way ANOVA with Tukey’s multiple comparisons correction. Scale bars are 2 µm.

Overall, Prex1 limits glucose clearance by limiting glucose uptake into liver cells, independently of its Rac-GEF activity. This requires the inhibitory action of Prex1 on the constitutively-active GPCR Gpr21, which enables Gpr21 to inhibit glucose uptake. Prex1-deficiency increases glucose uptake and mitochondrial glucose metabolism in liver cells. Mitochondrial size and membrane potential are increased in *Prex1^−/−^* liver cells, presumably to cope with the increased metabolism of excess glucose into ATP **(Graphical Abstract)**.

## Discussion

We show that Prex1 controls glucose homeostasis by maintaining fasting blood glucose levels and insulin sensitivity through its catalytic Rac-GEF activity, and by limiting glucose clearance into the liver through an adaptor function that involves the orphan GPCR Gpr21 and control of mitochondrial activity.

Prex1 deficiency protected mice on HFD from developing diabetes, without affecting body weight or body fat. Increased glucose clearance was seen in *Prex1^−/−^* mice throughout ageing and in both sexes, with some sex- and diet-related differences. The glucose tolerance of females was better than that of males throughout life, as expected (Mauvais-Jarvis et al., 2013; Reynolds et al., 2019), but glucose tolerance was improved in *Prex1^−/−^* males compared to *Prex^+/+^* more so on chow diet than HFD, whereas its was improved in *Prex1^−/−^* females mainly on HFD. The origin of these Prex1-dependent sex differences remains to be determined. The increased glucose tolerance was not merely a consequence of low fasting blood glucose, as the former was an adaptor function of Prex1, whereas the latter was Rac-GEF activity dependent. The insulin resistance of *Prex1^−/−^* mice was age-dependent, seen at ‘young-old’ (10 months) but not middle age (5 months), and stemmed mainly from an inability to maintain low blood glucose during the recovery phase after insulin administration, whereas the initial drop in blood glucose level was compromised less. This suggests an impaired counter-regulatory response to hypoglycaemia rather than *bona fide* insulin resistance (Alquier and Poitout, 2018; Hughey et al., 2014). In contrast to *Prex1^−/−^*, *Prex1^GD^* mice showed real insulin resistance. The difference may be due to our using lower dose insulin for *Prex1^GD^* mice, as their fasting blood glucose was lower, and we did not want to risk dangerous levels of hypoglycaemia. Alternatively, it could point to a dominant-negative role of catalytically-inactive Prex1, as the GEF-dead protein should retain the ability to bind PIP_3_ without being able to transmit Pi3k signals downstream. The mechanisms underlying the Rac-GEF activity dependent control of fasting blood glucose and insulin sensitivity remain to be elucidated. In this study, we focussed on the adaptor function of Prex1 in glucose clearance.

The increased glucose clearance in *Prex1^−/−^* mice was unexpected. Reports that Prex1 promotes glucose uptake into 3T3-L1 adipocytes (Balamatsias et al., 2011), and is required for the thermogenic capacity of brown adipocytes (Xue et al., 2015), predicted that Prex1 deficiency would impair glucose uptake and energy usage by adipocytes, and result in elevated blood glucose levels. Moreover, deficiency in the Prex1 homologue Prex2 causes glucose intolerance (Hodakoski et al., 2014), although we did not use this as a predictor, as Prex2 is thought to control glucose homeostasis through its inhibition of Pten (Hodakoski et al., 2014), which Prex1 does not do (Fine et al., 2009).

We pursued a number of avenues to identify the mechanisms underlying the altered glucose homeostasis. We verified that *Prex1^−/−^* mice have normal food and water consumption and waste production, and we and others previously showed them to have normal locomotor activity (Donald et al., 2008; Li et al., 2015). Histological analysis revealed no gross morphological abnormalities, and glucose excretion was normal, as were plasma levels of insulin, glucagon and adiponectin, and glucose-stimulated insulin secretion by pancreatic islets, the latter despite a report suggesting that P-Rex1 is required for insulin secretion in INS-1 832/13 pancreatic β cells (Thamilselvan et al., 2020). One exception was increased plasma glucagon upon insulin challenge, which might account in part for the inability of *Prex1^−/−^* mice to maintain low blood glucose after insulin administration and would merit future investigation. Both *Prex1^−/−^* and *Prex1^GD^* mice had small livers, so liver size should not be linked to the increased glucose clearance, which is seen only in *Prex1^−/−^* but not *Prex1^GD^* mice. Our analyses did not reveal obvious differences in hepatocyte size and numbers in *Prex1^−/−^* livers, so the origin of the small liver size remains to be determined.

Our signalling analysis revealed that human P-Rex1 promotes the insulin-stimulated activation of Akt upon overexpression in HepG2 cells, independently of its Rac-GEF activity, but endogenous Prex1 was not required for the insulin-stimulated activation of Akt, p70^S6K^, or GSK3β in metabolic tissues. However, Erk activity was constitutively increased in *Prex1^−/−^* but not *Prex1^GD^* liver cells, and GSK3β activity was increased in *Prex1^−/−^* adipose tissue upon insulin challenge. Knockdown of P-Rex1 in cancer cell lines is known to reduce the insulin receptor-dependent activation of Akt, Mek, and Rac1, as well as cell adhesion and proliferation (Dillon et al., 2015; Kim et al., 2011; Montero et al., 2013). It seems likely that endogenous Prex1 also mediates insulin signalling in primary liver, skeletal muscle, and adipose cells, but did not under the conditions investigated here. The constitutively increased Erk activity in liver cells was intriguing, as Erk is linked to increased fasting blood glucose and insulin resistance (Jiao et al., 2013), the opposite of what we see in *Prex1^−/−^* liver. Erk inhibition may shed light on the importance of Erk in the *Prex1^−/−^*glucose homeostasis phenotype.

Our analysis of glucose uptake revealed the liver as the critical organ through which Prex1 controls glucose homeostasis, with constitutively increased glucose uptake into *Prex1^−/−^* but not *Prex1^GD^*liver cells. The liver is the largest organ in the body, and the effect of Prex1 deficiency on hepatic glucose uptake was substantial, sufficient to explain the increased glucose clearance *in vivo*. Glucose uptake into skeletal muscle was also increased to some degree, whereas glucose uptake into various adipose tissues was normal. Presumably, the excess glucose enters *Prex1^−/−^* liver cells through the increased cell surface Glut2 and then undergoes increased metabolism to ATP in the mitochondria. This is aided by the constitutively increased mitochondrial size and membrane potential, which are likely a stress response of liver cells to the chronically increased glucose uptake in hepatocytes (Ke, 2020). It is not inconceivable, however, that the entire process might be driven by an adaptor function of Prex1 in limiting mitochondrial membrane potential, with the increased glucose uptake being a consequence of that. However, the identification of Gpr21 as a target of Prex1 in this process makes the latter possibility seem unlikely. We observed a large number of mega-mitochondria in *Prex1^−/−^* liver cells. These structures are associated with liver disease, including NAFLD and NASH (Shami et al, 2023). A recent report demonstrated increased Prex1 expression in the liver in human and mouse NAFLD, and used adenovirus-mediated knockdown of Prex1 in the mouse liver to show that Prex1 is required for the development of HFD-induced NAFLD (Li et al., 2022), supporting the importance of Prex1-dependent hepatic glucose homeostasis in diseases linked to metabolic syndrome. It is possible that the adenoviral treatment in that study may have had some off-target effects, as Prex1-depleted mice lost body weight, whereas the *Prex1^−/−^*mice used here did not. However, the study encourages the future investigation of adaptor functions Prex1 in other metabolic syndrome-related diseases of the liver, such as non-alcoholic steatohepatitis (NASH), cirrhosis, and liver cancer.

The only other known adaptor function of P-Rex1, apart from the control of glucose clearance, is its role in GPCR trafficking, which we recently described (Baker, Hampson et al., manuscript under review at Cell Reports). P-Rex1 inhibits the agonist-stimulated internalisation of all GPCRs tested, regardless of the type of heterotrimeric G protein the receptors couple to. In contrast, Prex1 does not control steady-state GPCR trafficking, nor the agonist-induced internalisation of receptor tyrosine kinases (Baker, Hampson et al., manuscript under review at Cell Reports). Furthermore, we know from previous work that Prex1 also does not affect the cell surface levels of integrins and selectins in neutrophils (Herter et al., 2013; Lawson et al., 2011; Pantarelli et al., 2021). Therefore, P-Rex1 plays a selective role in the agonist-induced trafficking of active GPCRs. Large portions of the P-Rex1 protein, including the PDZ, DEP and IP4P domains are required for this role. Mechanistically, P-Rex1 blocks the phosphorylation of active GPCRs at their C-terminal tail by Grk2, which is required for GPCR internalisation. P-Rex1 binds Grk2 directly, without obviously regulating Grk2 activity. We proposed that P-Rex1 limits the agonist-induced internalisation of GPCRs through its interaction with Grk2 to maintain high levels of active GPCR at the plasma membrane, thereby playing a dual role in promoting GPCR responses, by limiting the limiting the internalisation of GPCRs through an adaptor function as well as mediating GPCR signalling through its Rac-GEF activity.

We hypothesised that the control of hepatic glucose metabolism and of GPCR trafficking might be linked. As Prex1 deficiency reduces the levels of active GPCRs at the cell surface (Baker, Hampson et al., manuscript under review at Cell Reports), but glucose uptake and metabolism are increased in *Prex1^−/−^* liver cells, we searched for a candidate GPCR which inhibits these cell responses. This brought Gpr21 to our attention. As mentioned above, Gpr21 is an inhibitory, constitutively active Gα_q_-coupled orphan GPCR whose importance in glucose tolerance has been controversial (Gardner et al., 2012; Osborn et al., 2012; Wang et al., 2016), although this controversy seems resolved by a recent, independent *Gpr21^−/−^* mouse showing improved glucose tolerance (Riddy et al., 2021). We confirmed that knockdown of GPR21, or treatment with the inverse agonist GRA2, which bocks GPR21 activity, increases glucose uptake in HepG2 cells. We used cell fractionation to show that Prex1 controls the trafficking of Gpr21, as it does with other GPCRs (Baker, Hampson et al., manuscript under review at Cell Reports), with increased endosomal localisation of Gpr21 in *Prex1^−/−^*liver cells. GRA2-mediated blockade of Gpr21 activity increased glucose uptake and mitochondrial ATP production in *Prex^+/+^* and *Prex1^GD^* but not in *Prex1^−/−^* liver cells. Hence, Prex1 controls hepatic glucose uptake and metabolism through Gpr21, independently of its Rac-GEF activity. We propose that Prex1 does this by controlling the trafficking of Gpr21. Under the conditions tested, we found normal plasma membrane localisation of Gpr21 but increased endosomal localisation. It seems likely, that under conditions which permit receptor internalisation, Prex1 deficiency will reduce the cell surface level of Gpr21 by further promoting receptor endocytosis, thereby removing the Gpr21-mediated block of hepatic glucose uptake and metabolism. It remains to be seen if Prex1 also controls the surface level of Glut2 trafficking through Gpr21 or does so more directly. Gpr21 blockade by GRA2 did not affect Glut2 surface levels in *Prex^+/+^* liver cells, despite increasing glucose uptake and ATP production, whereas Glut2 surface levels were increased *Prex1^−/−^*cells. The P-Rex1 homologue P-Rex2 plays the same role in GPCR trafficking, so it would be worth revisiting glucose homeostasis in *Prex2^−/−^* mice, as our results suggest that P-Rex2 may regulate glucose homeostasis not entirely through inhibition of Pten, but also in part through GPCR trafficking.

Targeting of Prex1 seems beneficial to tackle metabolic disease, as Prex1 deficiency lowered fasting blood glucose, preventing mice on HFD from developing diabetic blood glucose levels. However, Prex1 deficiency also reduced insulin sensitivity, so any targeting of the catalytic activity of Prex1 would likely have this undesired effect. We recently developed the first inhibitors of P-Rex Rac-GEFs, which work on human and mouse P-Rex1, but these are laboratory tools with some cytotoxicity, not suitable for use *in vivo* (Lawson et al., 2022). In general, it is difficult to inhibit Rac-GEF activity, as Rac-GEFs operate through a relatively large surface interaction with Rac, so inhibitors often lack specificity and efficacy. However, GEF inhibition is not impossible, as shown by notable exception of brefeldin A, which inhibits Arf-GEFs and is widely used in cell biology to disassemble the Golgi complex (Cherfils and Zeghouf, 2013). Targeting of P-Rex1 by reducing P-Rex1 protein stability would have presumably the same undesirable effect of compromising insulin sensitivity. In comparison, targeting the adaptor function of Prex1 in glucose clearance seems a more promising avenue, as one would not have to target Prex1 itself, but could instead aim to inhibit Gpr21. The development of therapeutics based on the Gpr21 inverse agonist GRA2 which blocks the inhibitory action of Gpr21 on glucose uptake should have similarly beneficial effects on glucose clearance as Prex1 deficiency, without affecting the Rac-GEF activity dependent functions of Prex1.

## Acknowledgments

We thank the Babraham Gene Targeting Facility for their help with generating the *Prex1^GD^* mouse strain and the excellent staff of the Babraham Biological Support Unit for the passionate and expert care they take of our animals, and for help with experiments. We thank Chris Church from AstraZeneca for his expert advice on adipocytes, and John Findlay, Maynooth University, for his expertise on GPR21 and GRA2. ET received a UK Biotechnology and Biological Sciences Research Council (BBSRC) iCASE PhD studentship in collaboration with AstraZeneca. PAM received a PhD studentship from the BBSRC Doctoral Training Programme. The project was funded by Institute Strategic Programme Grant BB/P013384/1 from the BBSRC to the Babraham Institute Signalling Programme.

## Author contributions

JC, ET, PM, KM-G, AR, SC, AM, and HW designed, performed, and analysed experiments. EB, JS, GK, and DB designed and analysed experiments. DH and HW planned and supervised the project and procured funding. JC, ET and HW wrote the manuscript.

## STAR★Methods

### Key resources table

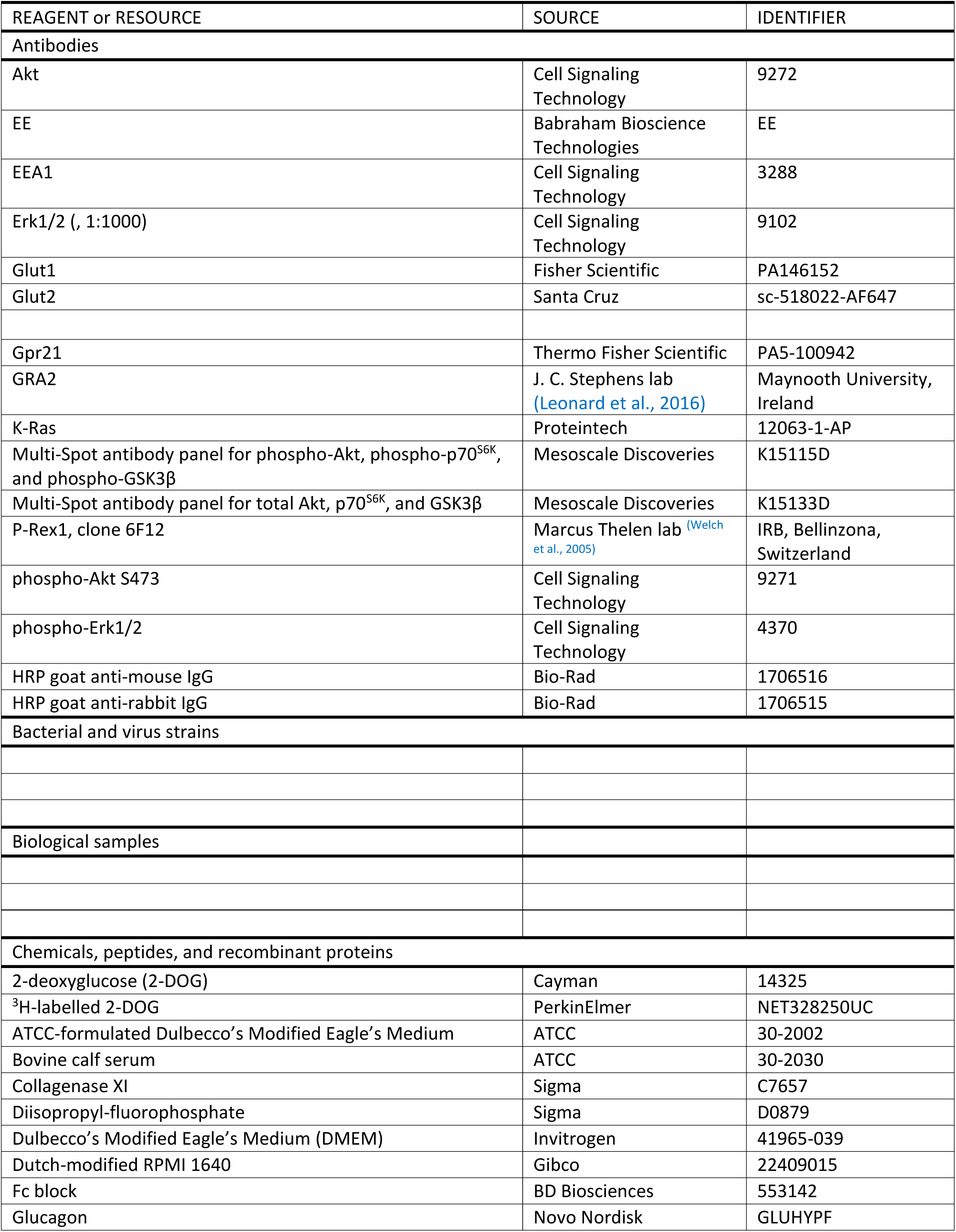
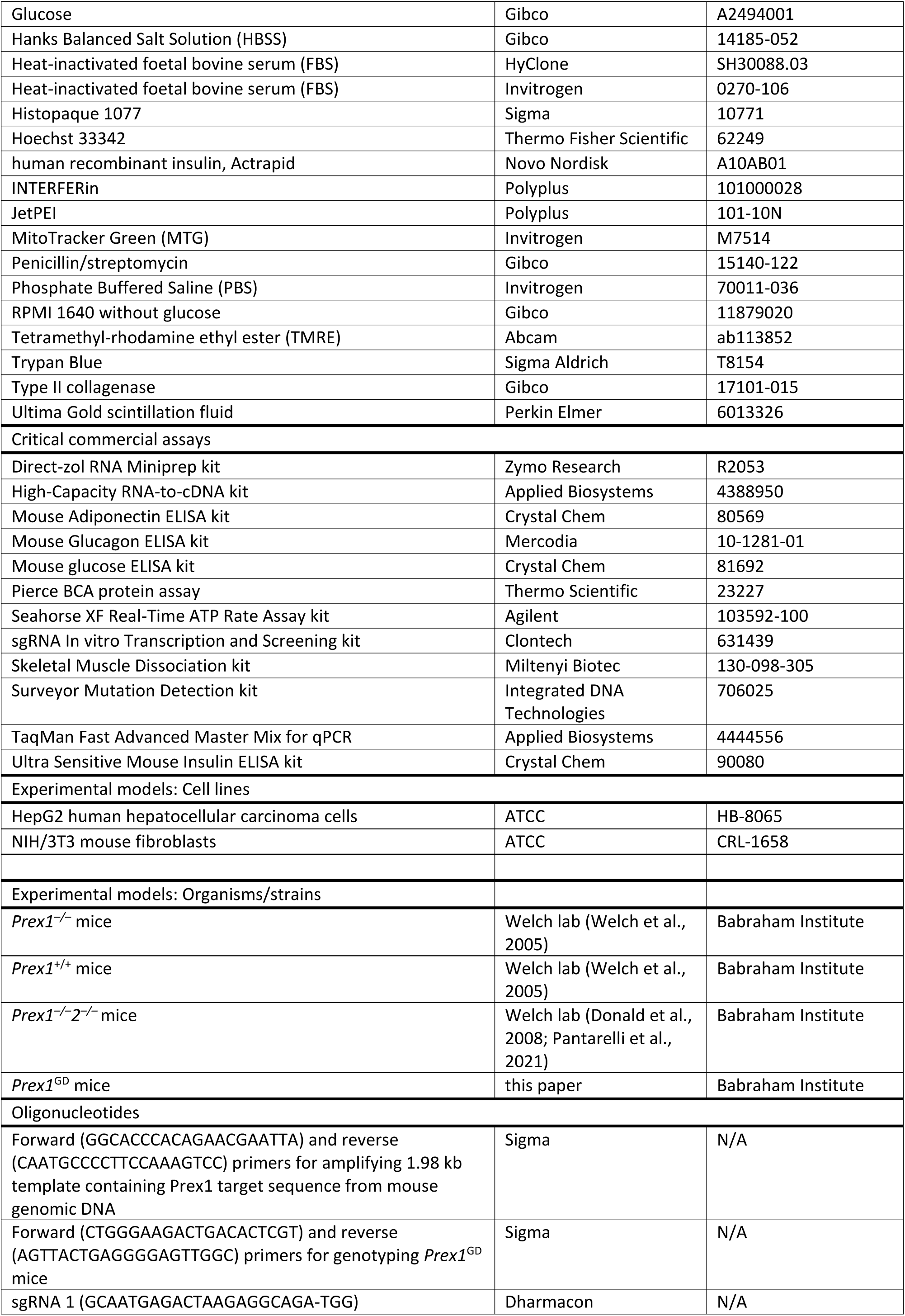
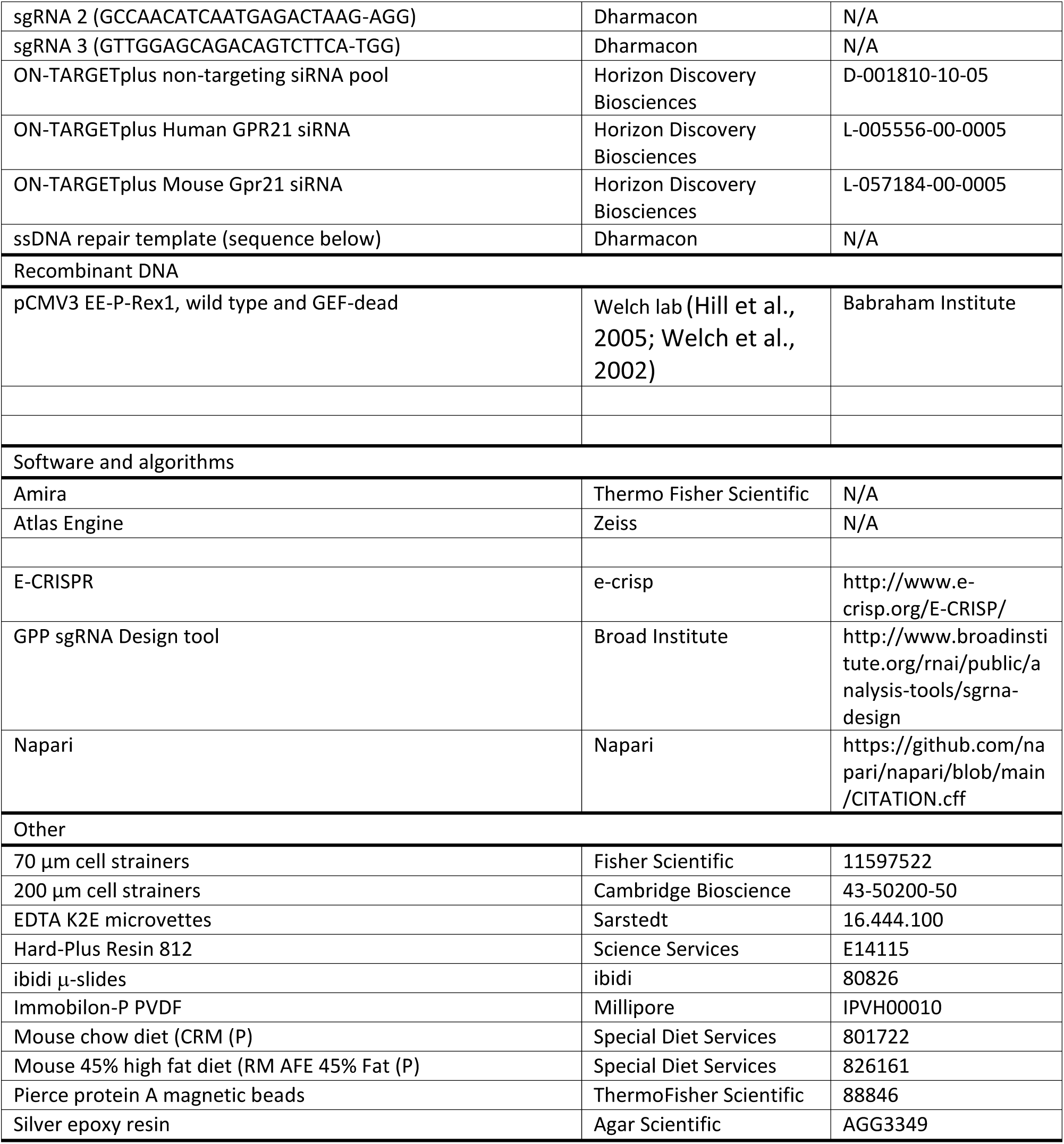

### Resource availability

#### Lead contact

Further information and requests for resources and reagents should be directed to and will be fulfilled by the lead contact, Heidi Welch.

#### Materials availability

Newly generated materials associated with the paper are available from the lead contact.

#### Data and code availability

- All data reported in this paper will be shared by the lead contact upon request.
- This paper does not report original code.
- Any additional information required to reanalyze the data reported in this work paper is available from the lead contact upon request.

### Experimental model and study participant details

Mouse strains and cell lines. Please see Key resources table and Method details sections ‘Mice’, ‘Generation of GEF-dead *Prex1^N233A, N233A^* mice’, and ‘Cell culture’.

### Method details

#### Mice

*Prex1*^−/−^ were previously described (Welch et al., 2005), backcrossed at least seven times to C57Bl6 genetic background, and compared to *Prex1*^+/+^ mice derived from the same backcrosses. To minimise genetic drift, *Prex1*^−/−^ and *Prex1*^+/+^ mice were intercrossed at least once every two years. *Prex1^−/−^2^−/−^* mice were as described (Donald et al., 2008; Pantarelli et al., 2021). Mice were group-housed (up to five) in individually ventilated cages in the Babraham Biological Support Unit which operates as described, using 12 h light/12 h dark cycles with dawn/dusk settings (Hornigold et al., 2022). For all experiments, mice were sex- and age-matched between genotypes. For *in vivo* experiments, they were additionally weight-matched at the onset. Prior to *in vivo* experiments, mice were habituated to handling by weighing, scruffing, and marking the tail with marker pen for rapid identification, for at least two days preceding the study. Mice were fed chow diet (CRM (P), Special Diets Services, Augy, France, 801722) *ad libitum* throughout, or were fed 45% HFD (RM AFE 45% Fat (P), Special Diets Services, 826161) *ad libitum* from the age of 10 weeks onwards. Unless specified otherwise, mice were tested at the age of 15 weeks. For ageing studies, at least two cohorts of 4-5 mice per genotype, sex, and diet were followed up to one year of age. Glucose tolerance was tested at 10, 15, 25, 39 and 52 weeks of age, insulin sensitivity at 25 and 43 weeks of age, and performance in metabolic cages at 11 months. At the end of the ageing study, mice were humanely killed and analysed by EchoMRI or dissected for organ analysis. Breeding and experiments were carried out with approval from the local Animal Welfare Ethical Review Body under the British Home Office Animal Scientific Procedures Act 1986.

#### Generation of GEF-dead Prex1^N233A, N233A^ mice

To generate a mouse strain with catalytically inactive (GEF-dead) Prex1, residue N233 in the catalytic DH domain was mutated to alanine using CRISPR/Cas gene editing. 20-nucleotide sgRNAs were designed to direct wild type Cas9 to the relevant site in exon 6, to be positioned directly upstream of a requisite 5’-NGG protospacer adjacent motif (PAM), and to have no highly homologous sites elsewhere in the mouse genome. Off-target potential was evaluated using software from the Zhang laboratory (http://crispr.mit.edu/), Broad Institute (http://www.broadinstitute.org/rnai/public/analysis-tools/sgrna-design), and E-CRISPR (http://www.e-crisp.org/E-CRISP/). Three sgRNAs were selected, sgRNA 1 (GCAATGAGACTAAGAGGCAGA-TGG), sgRNA 2 (GCCAACATCAATGAGACTAAG-AGG) and sgRNA 3 (GTTGGAGCAGACAGTCTTCA-TGG), the last 3 bps of each being the PAM. sgRNA efficiency was assessed using the sgRNA In vitro Transcription and Screening kit (Clontech, 631439) following the manufacturer’s protocol, with a 1.98 kb template containing the target sequence amplified from mouse genomic DNA using primers GGCACCCACAGAACGAATTA and CAATGCCCCTTCCAAAGTCC. All sgRNAs directed efficient cutting of the DNA by Cas9 *in vitro*. sgRNAs 1 and 3 were selected for further assessment of efficacy in cells by Surveyor Mutation Detection kit (Integrated DNA Technologies, 706025) following the manufacturer’s instructions. NIH/3T3 cells were transfected with pSpCas9(BB)-2A-GFP carrying the selected sgRNAs, genomic DNA was extracted, the relevant region was amplified, treated with Surveyor nuclease, and analysed by agarose gel electrophoresis. sgRNA 1 was chosen for targeting. A 199-nucleotide single-stranded ssDNA repair template was designed to introduce the desired point mutation by homology-directed repair, carry silent mutations creating restriction sites for screening purposes, and have symmetric homology arms of ≥90 nucleotides (ACTAGGAGTTGGCCAAGAGGACCCCCGGCAAGCACCCTGACCACACCGCGGTACAGAGTGCCCTGCAGGCC ATGAAGACTGTCTGCTCCAACATC**GCG**GAGAC**c**AAG**c**G**a**CA**a**ATGGAGAAGCTGGAGGCCCTGGAGCAGCTG CAGTCTCACATTGAAGGCTGGGAGGTGCGTGGTCAGGGTCCTCTGCTGAGGGGGCT). Nucleotides that introduce the N233A mutation are shown in bold capitals, silent mutations in bold lower case. The ssDNA repair template was purchased from Dharmacon as PAGE-purified Ultramer ssDNA.

The selected sgRNA, ssDNA repair template, and Cas9 mRNA were microinjected into C57BL/6J mouse zygotes by the Babraham Gene Targeting Facility. Pups were genotyped by sequencing 566 bp PCR products amplified using CTGGGAAGACTGACACTCGT forward and AGTTACTGAGGGGAGTTGGC reverse primers. Touch-down PCR was performed using Pfu, with denaturing at 95°C for 5 min; 3 cycles of 95°C, 30 s, annealing 30 s at 68°C, extension at 74°C for 2 min/kb; 3 cycles the same except annealing at 65°C; 3 cycles annealing 62°C; 35 cycles annealing at 59°C; and a final step at 74°C for 5 min. PCR products were isolated from agarose gels using QIAquick or GeneJET PCR purification kits and sequenced. Heterozygous *Prex1^N233A/+^*mice were crossed to C57BL/6J, then bred together to generate homozygous *Prex1^N233A/N233A^*animals. *Prex1^N233A/N233A^* mice were born at somewhat less than the expected Mendelian rate, were fertile, albeit with reduced litter sizes, and appeared healthy, with the white belly phenotype characteristic of *Prex1^−/−^* mice. Once the *Prex1^N233A/N233A^* (*Prex1*^GD^) strain was fully established, routine genotyping was done by Transnetyx (Cordova, TN, United States). *Prex1*^GD^ mice were compared to *Prex1*^+/+^ mice on the same C57BL/6J genetic background.

#### Cell culture

Mammalian cell lines were maintained at 37°C, 5% CO_2_, in a humidified atmosphere and used between passages 4 and 25. Aliquots of early-passage cells were stored in liquid N_2_. Human hepatocellular carcinoma HepG2 cells (ATCC, HB-8065) were cultured in flasks (Nunc) in Dulbecco’s Modified Eagle’s Medium (DMEM) (Invitrogen, 41965-039) supplemented with 10% heat-inactivated foetal bovine serum (FBS) (Invitrogen, 10270-106), unless stated otherwise, and were subcultured twice weekly using 0.25% (w/v) Trypsin, 0.53 mM EDTA. NIH/3T3 mouse fibroblasts (ATCC, CRL-1658) were cultured in flasks in ATCC-formulated DMEM (ATCC, 30-2002) with bovine calf serum (ATCC, 30-2030) and subcultured twice weekly using Trypsin/EDTA. For transient overexpression, transfection was done using JetPEI (Polyplus, 101-10N) following the manufacturer’s protocol. For knockdown experiments, HepG2 cells were cultured in DMEM, 10% heat-inactivated FBS (HyClone, SH30088.03), 100 U/ml penicillin and 100 μg/ml streptomycin (Gibco, 15140-122) for 6 h in a 6-well plate (5 × 10^4^ cells per well) in a humidified incubator at 37°C, 5% CO_2_. Cells were transfected with control (Horizon Discovery Biosciences, ON-TARGETplus non-targeting pool, D-001810-10-05) or GPR21 siRNA (Horizon Discovery Biosciences, ON-TARGETplus Human GPR21 siRNA, L-005556-00-0005), or ON-TARGETplus Mouse Gpr21 siRNA (L-057184-00-0005), as appropriate, using INTERFERin (Polyplus, 101000028) according to the manufacturer’s instructions, cultured for 72 h, and serum-starved in Krebs Ringer Buffer (KRB; 121 mM NaCl, 4.9 mM KCl, 1.2 mM MgSO_4_, 5 mM CaCl_2_, 12 mM Hepes) for 3 h prior to assays.

#### Glucose tolerance test

Mice were fasted for 6 h by moving into a fresh cage without food at 8.30 am, water remaining available *ad libitum*. 15 min prior to glucose injection, the mice were weighed and their baseline fasting blood glucose level was tested using an AlphaTRAK 2 glucometer (Abbott). At 2.30 pm, 2 g/kg glucose (20% glucose, Gibco, A2494001; 10 ml/kg) was administered by intraperitoneal (*i.p.*) injection. Tail bleeds were performed at 15, 30, 45, 60, 75 and 90 min after glucose injection and the blood glucose level determined by glucometer. For each timepoint, two glucometer readings were taken, and the lowest value recorded. Tail bleeds were done by pricking the tip of the tail (rather than the lateral tail vein) with a sterile needle for the first bleed, dislodging the scab for each following bleed, and without restraining the animal, to minimise stress and discomfort. Each bleed was no more than a few drops in blood volume. Mice on HFD at 39 and 52 weeks of age were injected with a lower dose of glucose (1 g/kg; 20% glucose at 5 ml/kg), to prevent dangerous levels of sustained hyperglycaemia.

#### Insulin sensitivity test

Mice were fasted for 4 h by moving into a fresh cage without food at 8.30 am, water remaining available *ad libitum*. 15 min prior to insulin injection, the mice were weighed, and their fasting blood glucose levels tested by glucometer as described here-above. At 12.30 pm, 0.75 IU/kg sterile human recombinant insulin (Actrapid, Novo Nordisk, A10AB01) in DPBS (5 ml/kg at 0.15 IU/ml) was administered by subcutaneous (*s.c.*) injection. Tail bleeds were performed at 15, 30, 45, 60, 90, 120 and 180 min after insulin injection and the blood glucose level determined by glucometer. For *Prex1^GD^*cohorts and their controls, a reduced insulin dose of 0.375 IU/kg was used. In rare cases, animals reached dangerously low blood glucose levels of 2-3 mM after insulin challenge. These mice were immediately injected *i.p.* with 100-150 μl 20% glucose to avoid hypoglycaemic shock, monitored, and excluded from the rest of the experiment.

#### Glucagon tolerance test

Mice were fasted for 6 h starting at 8.30 am, with water *ad libitum*, weighed, and their fasting blood glucose levels measured as described above. 5 ml/kg sterile glucagon (at 16 μg/kg; Novo Nordisk, GLUHYPF) was injected *i.p.*, and the blood glucose level determined by tail bleeds at 15, 30, 45, 60, 75, 90 min after the injection as described above.

#### Metabolic cages

To monitor food and water consumption and the production of urine and faeces, mice were group-housed (up to 3) in metabolic cages (Techniplast, UK), at the age of 11 months, unless otherwise indicated. Mice were habituated to the metabolic cages for 1 h on the first and 3 h on the second day before the experiment, then housed in the metabolic cages for 16 h overnight on 3 subsequent nights with pre-weighed food and water *ad libitum*. During daytime, the mice were returned to their home cages. The weights of urine and faeces and of the remaining food and water were recorded after each night. The metabolic cages were cleaned and stocked with fresh pre-weighed food and water before each testing night.

#### Body composition, organ weights and histology

At the end of ageing studies, 12 months-old mice were humanely killed by CO_2_ asphyxiation, death confirmed by pithing, carcasses stored at -80°C, thawed, and scanned by EchoMRI to determine body fat and lean mass. For organ weighing, mice of the indicated ages were humanely killed by CO_2_ asphyxiation (death confirmed by severing the femoral artery), weighed, and dissected for collection of brain, interscapular BAT, inguinal WAT, anterior thigh skeletal muscle, pancreas, kidneys, spleen, liver, thymus, heart, lung, salivary gland, and where applicable testes. The liver was scored visually for steatosis, from grade 0, with a healthy dark red colour, to grade 3, with a severe level of fat and yellow in colour overall. Organs were weighed and sometimes snap-frozen in liquid N_2_. For histological analysis, freshly dissected liver, skeletal muscle, inguinal WAT and interscapular BAT were fixed in formalin, and H&E sections were prepared by Abbey Veterinary Services and analysed by pathologist Dr Cheryl Scudamore, ExePathology, Exmouth, UK by blinded review using a grading system from 0 to 5, where 0 is normal and 5 is whole tissue affected.

#### Tissue and cell lysates, and western blotting

For preparation of tissue lysates, frozen organs were wrapped in cling film, shattered using a hammer, weighed, and lysed in 9 vol RIPA buffer (30 mM Hepes, pH 7.4, 150 mM NaCl, 1% Nonidet P-40, 0.5% deoxycholate, 0.1% SDS, 5 mM EGTA, 4 mM EDTA) supplemented with 1 mM DTT, 100 μM PMSF, and 10 μg/ml each of leupeptin, pepstatin-A, aprotinin and antipain (RIPA^+^). Samples were either sonicated in a Misonix S-3000 sonicator and incubated for 5 min on ice, or triturated frequently and incubated for 10 min on ice, and then centrifuged at 800 × g for 5 min at 4°C. Boiling 4× SDS-PAGE sample buffer was added to the supernatant (to final 1.3×), and samples were boiled for 5 min, snap frozen liquid N_2_, and stored at - 80°C. For adipose tissue, fat was removed first. The frozen tissue was shattered with a hammer, 500 μl isolation medium (50 mM Tris, 150 mM NaCl, 0.2 mM EDTA, 100 μM PMSF, and 10 μg/ml each of leupeptin, pepstatin-A, aprotinin and antipain) was added, followed by the addition of 1.875 ml CHCl_3_/MeOH, and incubation on ice for 15 min with sporadic mixing. 625 μl H_2_O was added, and samples were centrifuged at 800 × g for 5 min and 4°C. The protein disk at the interphase was collected, boiling 1.3× SDS-PAGE sample buffer added, and samples were boiled for 5 min, snap frozen, and stored at -80°C. To prepare lysates of cultured cells, cells were rinsed in ice-cold phosphate buffered saline (PBS, Invitrogen, 70011-036), scraped into RIPA^+^ buffer, samples were transferred into pre-cooled Eppendorf tubes, incubated on ice for 5 min, and debris was sedimented at 12,000 × g for 10 min at 4°C. Boiling 4× SDS-PAGE sample buffer was added to the supernatant to final 1.3×, and samples were boiled for 5 min, snap frozen, and stored at -80°C. To prepare lysates of mouse neutrophils, mature neutrophils were isolated from mouse bone marrow and treated with the cell-permeable serine protease inhibitor diisopropyl-fluorophosphate (Sigma, D0879) as described (Hornigold et al., 2022), before boiling 1.3× SDS-PAGE sample buffer added, and samples were boiled for 5 min, snap frozen, and stored at -80°C.

For western blotting, proteins were resolved by SDS-PAGE and blotted onto Immobilon-P membrane (Millipore, IPVH00010). Primary antibodies were EE (clone Glu-Glu, Babraham Bioscience Technologies, 1:50), P-Rex1 (clone 6F12, hybridoma supernatant, Thelen laboratory, Bellinzona, Switzerland, 1:30), GPR21 (Thermo Fisher Scientific, PA5-100942, 1:1000), Kras (Proteintech, 12063-1-AP, 1:6000) and the following antibodies from Cell Signaling Technology (London, UK), Eea1 (3288, 1:1000), phospho-Akt S473 (9271, 1:500), Akt (9272, 1:1000), phospho-Erk1/2 (4370, 1:2000), Erk1/2 (9102, 1:1000). Secondary antibodies were goat anti-mouse IgG-Horseradish Peroxidase [HRP] conjugate (BioRad, 1:3000) or goat anti-rabbit IgG-HRP (BioRad, 1:3000). Amersham enhanced chemiluminescence reagents ECL or ECLPlus were used for detection. To control for protein loading, gels or western blots were stained with 0.1% Coomassie Brilliant Blue R-250. Where required, blots were stripped in 25 mM glycine, pH 2.0, 1% SDS and reprobed. Western blots were quantified by densitometry using Fiji (ImageJ).

#### Liver enzymes

To quantify *Pepck* and *G6p* expression in the liver, mice were fasted for 4 h and then *s.c.* injected with either DPBS or 0.75 IU/kg insulin, humanely killed 15 min later by cervical dislocation, and the liver was excised rapidly and snap frozen in liquid N_2_. RNA was isolated using the Direct-zol RNA Miniprep kit (Zymo Research, R2053) following the manufacturer’s protocol. cDNA was synthesised using the High-Capacity RNA-to-cDNA kit (Applied Biosystems, 4388950) and qPCR performed using TaqMan Fast Advanced Master Mix (Applied Biosystems, 4444556) according to the manufacturer’s guidelines in a QuantStudio 12K Flex Real-Time PCR system.

#### Mesoscale analysis of insulin signalling

Mice were fasted, their fasting blood glucose levels tested, and injected *s.c.* with insulin as described for the insulin tolerance test, or mock-treated with sterile DPBS. 14 min after the injection, blood glucose levels were measured, and 15 min after the injection, the mice were humanely killed by cervical dislocation and death confirmed by pithing. Immediately, mice were dissected for collection of liver, anterior thigh skeletal muscle and inguinal WAT, and tissues were snap frozen in liquid N_2_ and stored at -80°C. Two biopsy punches of 2 mm were taken per tissue using disposable biopsy punches (KAI, BP-20F), and homogenised in Mesoscale Discoveries (MSD)-formulated lysis buffer (20 mM Tris, pH 7.5, 150 mM NaCl, 1 mM EDTA, 1 mM EGTA, 1% Triton-X-100, 1% protease inhibitors, 1% phosphatase inhibitors I and II, and 0.4% PMSF; MSD, R60TX), using a TissueLyser II (Qiagen) at setting 300 for 90 s. Samples were incubated for 30 min on ice with frequent vortexing, centrifuged at 1000 × g for 10 min at 4°C, and the supernatant was stored at -80°C. For skeletal muscle, an additional round of homogenisation was done prior to the addition of lysis buffer, and for adipose tissue a delipidation step was added after the centrifugation. Sample concentrations were determined by Pierce BCA protein assay (Thermo Scientific, 23227). 25 μl of tissue lysates (2125 μg/ml liver, 1500 μg/ml skeletal muscle, 800 μg/ml inguinal WAT) were tested using the Mesoscale Multi-Spot system to measure the phospho- and total levels of Akt, p70^S6K^, and GSK3β with antibody panels K15115D and K15133D in an MSD SECTOR imager.

#### Other signalling pathway analyses

To evaluate insulin signalling in HepG2 cells, cells were transfected with wild type or GEF-dead pCMV3-EE-P-Rex1 (Hill et al., 2005; Welch et al., 2002), or were mock-transfected, serum-starved for 8 h in serum-free medium containing 0.1% fatty-acid-free BSA, and stimulated with 25 nM insulin for various periods of time from 0 to 2 h. Cells were sedimented at 10,000 × g for 30 s at 4°C, and lysed in RIPA^+^ buffer for 5 min on ice with frequent vortexing. Debris was sedimented at 12,000 × g for 3 min at 4°C. Boiling 4× SDS-PAGE sample buffer was added to the supernatant (to final 1.3×), and samples were boiled for 5 min, snap frozen liquid N_2_, and stored at -80°C. Total cell lysates were western blotted for active Akt (phospho-S473). To control for the expression of wild type or GEF-dead P-Rex1, blots were probed with EE antibody.

To evaluate insulin and Gpr21 signalling in mouse liver cells, 1 × 10^6^ liver cells were stimulated with 100 nM insulin or treated with 30 μM GRA2, a small-molecule inverse agonist of Gpr21 (Leonard et al., 2016), for 10 min at 37°C. Ice-cold PBS was added, and cells were sedimented at 12,000 × g for 1 min at 4°C. Cells were lysed in RIPA^+^ buffer for 5 min on ice, debris was sedimented at 12,000 × g for 10 min at 4°C, boiling 4× SDS-PAGE sample buffer added to the supernatant to final 1.3×, and samples were boiled for 5 min, and then snap-frozen. Samples were analysed western blotting with phospho-Erk antibody (Cell Signalling Technology, 4370, 1:1000) before membranes were stripped and reprobed with total Erk antibody (Cell Signalling Technology, 9102, 1:1000).

#### ELISA

Mice were fasted from 8.30 am for 6 h and killed humanely by CO_2_ asphyxiation. Alternatively, they were fasted and challenged with glucose or insulin as described for the glucose and insulin tolerance tests, or mock-treated, and killed 15, 60 or 90 min later. Peripheral blood was collected immediately by cardiac puncture into 300 μl EDTA K2E microvettes (Sarstedt, 16.444.100). Samples were centrifuged for 5 min at 2000 × g at RT, and the supernatant was transferred into fresh tubes and centrifuged again. The second supernatant (plasma) was aliquoted and stored at -80°C. To quantify plasma adiponectin, glucagon, and insulin, the Mouse Adiponectin (Crystal Chem, 80569), Glucagon (Mercodia, 10-1281-01), and Ultra Sensitive Mouse Insulin (Crystal Chem, 90080) ELISA kits were used, following manufacturers’ guidelines, measuring A_450_ on a PHERAstar FS microplate reader. To analyse glucose in urine, mice were fasted and challenged with glucose as described for the glucose tolerance test, scruffed 90 min after glucose injection, and urine was collected into an Eppendorf tube. A mouse glucose ELISA (CrystalChem, 81692) was used according to the supplier’s protocol using measurements at A_505_.

#### Insulin release from isolated pancreatic islets

15-week-old mice were humanely killed by cervical dislocation and death confirmed by pithing. The pancreas was recovered, injected with 5 ml ice-cold collagenase XI (1000 U/ml, Sigma, C7657) in HBSS (Gibco, 14185-052) with 1 mM CaCl_2_, and placed into a 50 ml falcon tube which was swirled in a 37°C water bath for 20 min, with shaking every 3 min for 1 min. Ice-cold HBSS with 1 mM CaCl_2_ was added to give a volume of 30 ml, and samples were centrifuged at 326 × g for 2 min. Samples were washed again, resuspended in 5 ml HBSS with 1 mM CaCl_2_, slowly added on top of 7.5 ml Histopaque 1077 (Sigma, 10771), centrifuged at 2000 × g for 15 min, brake 0, and the middle layer containing islets was transferred into a 5 cm dish.

20 islets were hand-picked under a dissection microscope using a sterile wide-neck 1 ml plastic pipette, placed into each well of a 6-well plate containing Dutch-modified RPMI 1640 (Gibco, 22409015) with 10% FBS, 1% L-glutamine, 100 U/ml penicillin and 100 μg/ml streptomycin, and left to recover ON in a humidified 37°C, 5% CO_2_ incubator. 5 or 10 islets, as stated, were handpicked into 2 ml Eppendorf tubes containing 1 ml RPMI 1640 without glucose (Gibco, 11879020), the volume adjusted to 2 ml, and samples were left to recover for 1 h in a humidified 37°C, 5% CO_2_ incubator, before being placed into a thermomixer at 37°C without shaking. A 50 µl aliquot was removed, taking care not to dislodge the islets resting at the bottom, and snap-frozen in liquid N_2_. 50 µl RPMI 1640 containing 80 mM glucose was added (for 2 mM final), and tubes were gently inverted twice to ensure mixing and incubated for 15 min. After 14 min, the tubes were inverted again, and at 15 min, once the islets had sunk to the bottom, another aliquot was taken. 50 µl fresh media containing 720 mM glucose (for 20 mM final) was added for a further 45 min, after which a final aliquot was taken. Secreted insulin was quantified by ELISA as described here-above.

To determine the total amount of insulin in pancreatic islets, 5 islets were sedimented for 5 min at 326 × g at 4°C and lysed in 1 ml ice-cold RIPA^+^ buffer by incubation on ice for 5 min with frequent vortexing. Samples were centrifuged at 10,000 × g for 3 min at 4°C, and the supernatant was snap-frozen in liquid N_2_. These samples were diluted 1:10 before analysis by insulin ELISA.

#### Isolation of primary liver cells, adipocytes, and skeletal muscle cells

To isolate liver cells, mice were culled by cervical dislocation, death was confirmed by pithing, and the liver perfused with PBS by cannulating the hepatic portal vein using a 25G needle, followed by cutting the inferior vena cava to drain blood and perfusion fluid. The perfused liver was quickly excised, minced using scissors, rinsed in DMEM for 30 min at RT with gentle agitation, passed through a 70 μm cell strainer (Fisher Scientific, 11597522), and rinsed with 5 ml DMEM. Cells were sedimented at 100 × g for 3 min at RT, resuspended in KRB, 0.1% BSA and incubated at 37°C for 90 min prior to use. Liver cells were counted by haemocytometer and viability assessed by Trypan Blue (Sigma Aldrich, T8154) exclusion. For Seahorse assays (see below), liver cells were resuspended in DMEM, 10% FBS, 100 U/ml penicillin and 100 μg/ml streptomycin.

To isolate adipocytes, mice were humanly killed by CO_2_ asphyxiation, death was confirmed by severing the femoral artery, and mouse adipose tissues (BAT, visceral WAT and subcutaneous WAT) were carefully excised, minced using scissors, and digested in a 50 ml Falcon tube containing 10-15 ml digestion buffer (KRB with 3% BSA, 1mg/ml type II collagenase (Gibco, 17101-015), 4 mM glucose, and 500 nM adenosine) for 1 h at 37°C, with gentle inverting every 10 min. EDTA was added to final 2 mM, and the homogenate was passed through a 200 μm cell strainer (Cambridge Bioscience, 43-50200-50). To collect mature adipocytes specifically, mature adipocytes were allowed to float to the top of the filtered homogenate for 1-2 h at RT, and the floating white cell layer was carefully harvested using a plastic Pasteur pipette. Adipocytes were counted using a haemocytometer, resuspended in KRB, 0.1% BSA, and incubated in a 12-well plate for 3 h at 37°C, 5% CO_2_ prior to use.

To isolate skeletal muscle cells, 1.5 g medial gastrocnemius, semimembranosus, and rectus femoris muscles were collected from mouse hind limbs, minced into 2-4 mm pieces using scissors, and dissociated by incubation with Skeletal Muscle Dissociation Kit enzymes (Miltenyi Biotec, 130-098-305) according to the manufacturer’s instructions for 30 min at 37°C in a GentleMACS C Tube under continuous gentle rotation, followed by homogenisation in a GentleMACS Dissociator (Miltenyi). The skeletal muscle cells were filtered through a 70 μm cell strainer, sedimented at 300 × g for 20 min, resuspended in KRB, 0.1% BSA, and incubated for 90 min at 37°C prior to stimulation.

#### Glucose uptake

1 × 10^6^ liver cells in 200 μl KRB, 0.1% BSA in Eppendorf tubes were stimulated with 200 μl 2× insulin (Actrapid, 100 nM final) for 10 min at 37°C, or were mock-stimulated, followed by the addition of 200 μl of 3× 2-deoxyglucose (2-DOG; Cayman, 14325, for 50 μM final) and 0.25 μCi ^3^H-labelled 2-DOG (PerkinElmer, NET328250UC) and further incubation for 60 min. Alternatively, liver cells were treated with 30 μM GRA2, for 3 h at 37°C, or were mock-treated, before further treatment as here-above. Liver cells were washed three times in ice-cold KRB and lysed in RIPA buffer for 5 min on ice. Lysates were mixed with Ultima Gold scintillation fluid (Perkin Elmer, 6013326), and glucose uptake was measured by scintillation counting (Packard 1600TR, Liquid Scintillation Analyser).

Mature adipocytes (isolated from either visceral or subcutaneous white, or from brown adipose tissue) in a 12-well plate 1x 10^6^ per well in 1 ml KRB, 0.1% BSA were stimulated with 250 μl 5× insulin (for 100 nM final) for 10 min at 37°C, or were mock-stimulated, followed by the addition of 250 μl 6× 2-DOG (for 50 μM final) and 0.25 μCi ^3^H-2-DOG, and further incubation for 60 min. 1 ml liquid was carefully aspirated from the bottom of the well, and 1.5 ml ice-cold KRB added to the floating cells. 1.5 ml of liquid was again removed and replaced with fresh buffer, and this wash was repeated 5 times. After the final aspiration, 250 μl RIPA buffer was added, and samples were incubated on ice for 5 min and vortexed hard. Glucose uptake was measured as described for liver cells.

200 μl skeletal muscle cells were stimulated with 100 nM insulin for 10 min at 37°C in Eppendorf tubes, or were mock-stimulated, followed by the addition of 50 μM 2-DOG and 0.25 μCi ^3^H-2-DOG, and further incubation for 60 min. Cells were washed three times in ice-cold KRB by centrifugation at 1500 × g for 3 min at 4°C, and lysed in RIPA buffer for 5 min on ice. Glucose uptake was measured as measured as described above.

To measure glucose uptake in transfected HepG2 cells, the cell culture medium was replaced with KRB, 0.1% BSA for 90 min, and cells were treated with 30 μM GRA2 for 3 h at 37°C, or were mock-treated. 50 μM 2-DOG and 0.25 μCi ^3^H-2-DOG were added for a further 60 min before ice-cold KRB was added to terminate the reaction. Cells were scraped into 250 μl ice-cold RIPA buffer, lysed for 5 min on ice, and debris was sedimented at 10 000 × g for 1 min at 4°C. Glucose uptake was measured as described above.

#### Cell surface level of glucose transporters

To evaluate the cell surface levels of glucose transporters, 1 × 10^6^ liver cells in 200 μl KRB, 0.1% BSA were stimulated with 5 mM glucose or 100 nM insulin for 10 min at 37°C, or were mock-stimulated, and sedimented at 300 × g for 5 min at 4°C. Alternatively, cells were stimulated with 100 nM insulin for 10 min at 37°C cells, and 5 mM glucose was added for another 30 min, or cells were treated with 30 μM GRA2 for 3 h at 37°C, or mock-treated, and sedimented. Cells were stained with antibodies to Glut2 (Santa Cruz, sc-518022-AF647, 1:200) or Glut1 (Fisher Scientific, PA146152, 1:200) in the presence of Fc block (BD Biosciences, 553142, 1:1000) for 30 min on ice in the dark, washed three times with ice-cold PBS, and resuspended in 200 μl PBS. Flow cytometry was performed using a BioRad ZE5 flow cytometer, recording 20,000 events per sample. The mean fluorescence intensity (mfi) of Glut2 and Glut1 signal was quantified using FlowJo. To determine total cellular levels of the glucose transporters, cells were permeabilised in 0.1% Triton X-100/PBS for 10 min on ice and washed twice in PBS prior to staining.

#### ATP production (Seahorse)

ATP production was analysed by measuring oxygen consumption rate (OCR) and extracellular acidification rate (EAR) in an Agilent Seahorse XF96 analyser by real-time ATP rate assay (103592-100), according to the manufacturer’s protocol. Unless otherwise stated, all reagents were from Agilent Technologies. In brief, 200 μl liver cells in DMEM, 10% FBS, 100 U/ml penicillin and 100 μg/ml streptomycin were plated at a density of 2 x 10^4^/well in a poly-D-lysine (0.1 mg/ml)-coated 96-well plate (101085-004) and incubated ON in a humidified incubator at 37°C, 5% CO_2._ A sensor cartridge was hydrated overnight at 37°C. Cells were rinsed and incubated with warmed assay medium (DMEM [103575-100, pH 7.4], 10 mM glucose [103577-100], 1 mM pyruvate [103578-100], 2 mM glutamine [103579-100]) at 37°C for 45-60 min prior to the assay. 1.5 μM oligomycin and 0.5 μM rotenone/antimycin A were loaded into the sensor cartridge according to the manufacturer’s instructions for induced ATP-rate assay. The Seahorse analyser was run using the standard initial equilibration step, then treatment with 100 nM insulin or 30 μM GRA2 for 2 h, with readings taken every 10 min, followed by the addition of oligomycin for 18 min and then rotenone for another 18 min, with readings taken every 6 min. ATP production from glycolysis and mitochondrial respiration was analysed using WAVE software (Agilent Technologies).

#### Liver cell fractionation

2 × 10^7^ liver cells were resuspended in 1.1 ml ice-cold, detergent-free lysis buffer (10 mM Pipes, pH 6.0 (on ice), 100 mM KCl, 3 mM NaCl, 3.5 mM MgCl_2_, 1 mM EDTA, 1 mM ATP, 25 µg/ml leupeptin, 25 µg/ml pepstatin A, 25 µg/ml aprotinin, 25 µg/ml antipain), and were lysed by gentle sonication in a Misonix Ultrasonicator XL with two pulses of 2 s on setting 3 (∼4 Watts), with 28 s breaks on ice for cooling. We verified by haematocytometer that this procedure was gentle enough to leave 5-10% of cells intact. 0.5 M EGTA was added to final 6 mM, and samples were centrifuged at 1000 × g for 10 min at 4°C to sediment live cells, nuclei and debris. 1 ml of the postnuclear supernatant was transferred into fresh pre-cooled 1.5 ml tubes and centrifuged at 10000 × g for 15 min to remove heavy organelles. 950 μl of the supernatant were transferred into a fresh, precooled tube, and whole endosomes were immunoprecipitated as described (Gosney, 2016), by incubation with EEA1 antibody (Cell Signaling technology 3288, 1:1000) for 1 h on ice with end-over-end rotation. Prewashed Pierce protein A magnetic beads (ThermoFisher Scientific, 88846) were added, and samples were incubated for a further 40 min. The beads containing immunoprecipitated endosomes were pelleted using magnets and washed 3× in lysis buffer. The post-immunoprecipitation supernatant was transferred into precooled ultracentrifuge tubes and ultracentrifuged in an MLT130 rotor at 200,000 × g for 30 min at 4°C. The plasma membrane pellet was washed once by careful addition and aspiration of 100 μl ice-cold lysis buffer. Boiling 1.3× SDS-PAGE sample buffer was added to both endosomal and plasma membrane fractions, and samples were boiled for 5 min and then snap frozen in liquid N_2_. Samples were analysed by western blotting with Gpr21, Eea1, and Kras antibodies.

#### Mitochondrial membrane potential

To evaluate mitochondrial membrane potential by flow cytometry, 1 × 10^6^ liver cells in 200 μl KRB, 0.1% BSA were stimulated with 5 mM glucose or 100 nM insulin, or were treated with 30 μM GRA2 for 30 min at 37°C, or mock-treated. Ice-cold PBS was added, and the cells were sedimented at 300 × g for 5 min at 4°C, stained with cell-permeable MitoTracker Green (MTG, Invitrogen, M7514, 100 nM) and tetramethyl-rhodamine ethyl ester (TMRE, Abcam, ab113852, 200 nM) dyes for 30 min on ice in dark, and washed three times with ice-cold PBS. Flow cytometry was performed using a BioRad ZE5 flow cytometer, recording 20,000 events per sample. The mfi of MitoTracker Green and TMRE was quantified using FlowJo.

To evaluate mitochondrial membrane potential by immunofluorescence microscopy, 200 μl liver cells in DMEM were plated into poly-D-lysine (0.1 mg/ml)-coated ibidi μ-slides (ibidi, 80826) at a density of 2 x 10^6^/well for 2 h in a humidified incubator at 37°C, 5% CO_2_. The medium was removed, and cells were stimulated with 5 mM glucose or 100 nM insulin in prewarmed DMEM for 30 min at 37°C, or were mock-stimulated. Alternatively, cells were treated with 30 μM GRA2 for 2 h at 37°C, or mock-treated. 15 min before the end of incubation, MTG, TMRE, and the cell permeable DNA dye Hoechst 33342 (Thermo Fisher Scientific, 62294, 1:400) were added, and the cells were live-imaged using a Nikon Eclipse Ti-E widefield system. Images were analysed by Fiji software for MTG and TMRE signal per field of view (fov).

#### Electron microscopy

To prepare liver samples for focussed ion beam scanning electron microscopy (FIB-SEM), liver tissue was excised and fixed in 2.5% glutaraldehyde and 4% formaldehyde in 0.2 M phosphate buffer for 30 min at RT, trimmed into 1-2 mm cubes, and fixed for a further 60 min. The fixed tissue was prepared according to the Ellisman protocol (https://www.protocols.io/view/preparation-of-biological-tissues-for-serial-block-36wgq7je5vk5/v2). Briefly, the tissue was washed in 0.1 M phosphate buffer followed by incubation in 2% osmium tetroxide containing 1.5% potassium ferrocyanide in 2 M phosphate buffer for 60 min. Samples were washed in distilled water and incubated in 1% thiocarbohydrazide for 20 min, washed in distilled water, incubated in 2% osmium tetroxide for 30 min, washed in distilled water, and placed in 2% uranyl acetate ON at 4°C. The following day, samples were stained *en bloc* with Walton’s lead aspartate for 30 min at 60°C followed by washing in distilled water and dehydration in graded ethanol and then propylene oxide. The tissue was embedded in Hard-Plus Resin 812 (Science Services, E14115), mounted on SEM stubs using silver epoxy resin (Agar Scientific, AGG3349). Liver tissue was revealed using a Leica UC6 microtome (Leica, Vienna, Austria), then coated with a 5 nm layer of platinum in a CU10 coater (Safematic, Switzerland).

For FIB-SEM, specimens were placed into a Zeiss Crossbeam 550 scanning electron microscope running Atlas Engine (Zeiss, Oberkoch, Germany). A region of interest was located in the SEM by imaging at 10 kV. The area was prepared for imaging by depositing tracking marks and milling a trench with the FIB beam to allow SEM imaging of the milled face. Electron micrographs were obtained with isotropic 10 nm resolution with a 9 or 10 µs dwell time. The SEM was operated at 2kV accelerating voltage and 500 pA current. Both backscatter and secondary electron data were collected, to improve signal to noise ratio. Ion beam milling was performed at an accelerating voltage of 30 kV and current of 700 pA.

For image analysis, image stacks were initially aligned using Atlas 5 software, before the combined image channels were exported. Aligned images were exported into Amira (Thermo Fisher Scientific, Waltham, MA, USA) and cropped to a region of interest. 800 voxels^3^ were extracted from the data set, corrected for stage tilt, and denoised using the dual-beam wizard. The corrected volume was exported to Napari (https://github.com/napari/napari/blob/main/CITATION.cff) (napari contributors, 2019) and opened using the Empanada (EM Panoptic Any Dimension Annotation) plugin. A training data set consisting of 16 patches of 256 pixels from a *Prex1^−/−^* data set were used to create and fine-tune a panoptic segmentation model, based on MitoNet (Conrad and Narayan, 2023). The refined model (MENL-v2) generated automated semantic segmentation of ER and nucleus and instance segmentation of lipid drops and mitochondria. Feature label files were imported to Amira for clean-up, segmentation, volume rendering and analysis. Segmented mitochondrial volumes were separated (using the separate objects function), labelled, and mitochondrial number, volume, and morphometrics and occurrence of mega-mitochondria were quantified using the ‘analyse labels’ function.

### Quantification and statistical analysis

Sample size was determined using power calculations to yield 80% power, based on results of pilot experiments and on previously published data as referenced. Experiments were performed at least three times except where indicated. Sample size and numbers of independent experiments are detailed in figure legends. Animals for experimental cohorts was selected as described under ‘Mice’ according to genotype, group size, sex, age, and weight. Within these criteria, mice were selected for cohorts at random by the staff of the Biological Support Unit. Image analysis was performed in a blinded manner. Data were tested for normality of distribution to determine if parametric or non-parametric methods of analysis were appropriate. Where warranted, data were log-transformed or square-root transformed prior to statistical analysis. For comparison of two groups, unpaired or paired Student’s t-test was used, as appropriate, whereas for comparison of multiple groups, one-way or two-way ANOVA was used, as appropriate, with repeated measures followed by post-hoc test with multiple comparisons correction. Data in categories were analysed by chi-square test. Where data were normalised, statistical analysis was done on raw data, prior to normalisation. Parameters with values of p ≤ 0.05 were considered to differ significantly. In the figures, p-values in black denote significant differences, p-values in grey are non-significant. In time courses, * indicates p < 0.05, ** p < 0.01, *** p < 0.001, and **** p < 0.0001. Results are presented as mean ± standard error of the mean (SEM). The number of experimental repeats is indicated in the figure legends. Statistical analysis and plotting of graphs were performed in GraphPad Prism 10.

## References

Alquier, T., and Poitout, V. (2018). Considerations and guidelines for mouse metabolic phenotyping in diabetes research. Diabetologia 61, 526–538.

Balamatsias, D., Kong, A.M., Waters, J.E., Sriratana, A., Gurung, R., Bailey, C.G., Rasko, J.E., Tiganis, T., Macaulay, S.L., and Mitchell, C.A. (2011). Identification of P-Rex1 as a novel Rac1-guanine nucleotide exchange factor (GEF) that promotes actin remodeling and GLUT4 protein trafficking in adipocytes. J Biol Chem 286, 43229–43240.

Bento, J.L., Palmer, N.D., Zhong, M., Roh, B., Lewis, J.P., Wing, M.R., Pandya, H., Freedman, B.I., Langefeld, C.D., Rich, S.S., et al. (2008). Heterogeneity in gene loci associated with type 2 diabetes on human chromosome 20q13.1. Genomics 92, 226–234.

Cherfils, J., and Zeghouf, M. (2013). Regulation of small GTPases by GEFs, GAPs, and GDIs. Physiol Rev 93, 269–309.

Conrad, R., and Narayan, K. (2023). Instance segmentation of mitochondria in electron microscopy images with a generalist deep learning model trained on a diverse dataset. Cell Syst 14, 58–71 e55.

Dillon, L.M., Bean, J.R., Yang, W., Shee, K., Symonds, L.K., Balko, J.M., McDonald, W.H., Liu, S., Gonzalez-Angulo, A.M., Mills, G.B., et al. (2015). P-REX1 creates a positive feedback loop to activate growth factor receptor, PI3K/AKT and MEK/ERK signaling in breast cancer. Oncogene 34, 3968-3976.

Donald, S., Humby, T., Fyfe, I., Segonds-Pichon, A., Walker, S.A., Andrews, S.R., Coadwell, W.J., Emson, P., Wilkinson, L.S., and Welch, H.C.E. (2008). P-Rex2 regulates Purkinje cell dendrite morphology and motor coordination. Proc Natl Acad Sci U S A 105, 4483–4488.

Dutta, S., and Sengupta, P. (2016). Men and mice: Relating their ages. Life Sci 152, 244–248.

Fine, B., Hodakoski, C., Koujak, S., Su, T., Saal, L.H., Maurer, M., Hopkins, B., Keniry, M., Sulis, M.L., Mense, S., et al. (2009). Activation of the PI3K pathway in cancer through inhibition of PTEN by exchange factor P-Rex2a. Science 325, 1261–1265.

Gardner, J., Wu, S., Ling, L., Danao, J., Li, Y., Yeh, W.C., Tian, H., and Baribault, H. (2012). G-protein-coupled receptor GPR21 knockout mice display improved glucose tolerance and increased insulin response. Biochem Biophys Res Commun 418, 1–5.

Gerich, J.E. (2000). Physiology of glucose homeostasis. Diabetes Obes Metab 2, 345–350.

Ghalali, A., Wiklund, F., Zheng, H., Stenius, U., and Hogberg, J. (2014). Atorvastatin prevents ATP-driven invasiveness via P2X7 and EHBP1 signaling in PTEN-expressing prostate cancer cells. Carcinogenesis 35, 1547–1555.

Gosney, J.A. (2016) Isolation of EGFR-containing early endosomes. PhD thesis. University of Louisville.

Hernandez-Negrete, I., Carretero-Ortega, J., Rosenfeldt, H., Hernandez-Garcia, R., Calderon-Salinas, J.V., Reyes-Cruz, G., Gutkind, J.S., and Vazquez-Prado, J. (2007). P-Rex1 links mammalian target of rapamycin signaling to Rac activation and cell migration. J. Biol. Chem. 282, 23708–23715.

Herter, J.M., Rossaint, J., Block, H., Welch, H., and Zarbock, A. (2013). Integrin activation by P-Rex1 is required for selectin-mediated slow leukocyte rolling and intravascular crawling. Blood 121, 2301–2310.

Hill, K., Krugmann, S., Andrews, S.R., Coadwell, W.J., Finan, P., Welch, H.C., Hawkins, P.T., and Stephens, L.R. (2005). Regulation of P-Rex1 by phosphatidylinositol (3,4,5)-trisphosphate and Gβγ subunits. J Biol Chem 280, 4166-4173.

Hodakoski, C., Hopkins, B.D., Barrows, D., Mense, S.M., Keniry, M., Anderson, K.E., Kern, P.A., Hawkins, P.T., Stephens, L.R., and Parsons, R. (2014). Regulation of PTEN inhibition by the pleckstrin homology domain of P-REX2 during insulin signaling and glucose homeostasis. Proc Natl Acad Sci U S A 111, 155–160.

Hornigold, K., Chu, J.Y., Chetwynd, S.A., Machin, P.A., Crossland, L., Pantarelli, C., Anderson, K.E., Hawkins, P.T., Segonds-Pichon, A., Oxley, D., et al. (2022). Age-related decline in the resistance of mice to bacterial infection and in LPS/TLR4 pathway-dependent neutrophil responses. Front Immunol 13, 888415.

Hornigold K, T.E., Pantarelli C, Welch HCE (2018; ). P-Rex1. In Encyclopedia of Signaling Molecules. S. Choi, ed., pp. 4142-4154.

Hughey, C.C., Wasserman, D.H., Lee-Young, R.S., and Lantier, L. (2014). Approach to assessing determinants of glucose homeostasis in the conscious mouse. Mamm Genome 25, 522–538.

Jiao, P., Feng, B., Li, Y., He, Q., and Xu, H. (2013). Hepatic ERK activity plays a role in energy metabolism. Mol Cell Endocrinol 375, 157–166.

Ke, P.Y. (2020). Mitophagy in the pathogenesis of liver diseases. Cells 9.

Kim, E.K., Yun, S.J., Ha, J.M., Kim, Y.W., Jin, I.H., Yun, J., Shin, H.K., Song, S.H., Kim, J.H., Lee, J.S., et al. (2011). Selective activation of Akt1 by mammalian target of rapamycin complex 2 regulates cancer cell migration, invasion, and metastasis. Oncogene 30, 2954–2963.

Kinsella, G.K., Cannito, S., Bordano, V., Stephens, J.C., Rosa, A.C., Miglio, G., Guaschino, V., Iannaccone, V., Findlay, J.B.C., and Benetti, E. (2021). GPR21 inhibition increases glucose-uptake in HepG2 cells. Int J Mol Sci 22.

Lawson, C., Donald, S., Anderson, K., Patton, D., and Welch, H. (2011). P-Rex1 and Vav1 cooperate in the regulation of fMLF-dependent neutrophil responses J Immunol 186, 1467-1476.

Lawson, C.D., Hornigold, K., Pan, D., Niewczas, I., Andrews, S., Clark, J., and Welch, H. (2022). Small-molecule inhibitors of P-Rex guanine-nucleotide exchange factors. Small GTPases 13, 307–326.

Leonard, S., Kinsella, G.K., Benetti, E., and Findlay, J.B.C. (2016). Regulating the effects of GPR21, a novel target for type 2 diabetes. Scientific reports 6, 27002.

Lewis, J.P., Palmer, N.D., Ellington, J.B., Divers, J., Ng, M.C., Lu, L., Langefeld, C.D., Freedman, B.I., and Bowden, D.W. (2010). Analysis of candidate genes on chromosome 20q12-13.1 reveals evidence for BMI mediated association of PREX1 with type 2 diabetes in European Americans. Genomics 96, 211–219.

Li, J., Chai, A., Wang, L., Ma, Y., Wu, Z., Yu, H., Mei, L., Lu, L., Zhang, C., Yue, W., et al. (2015). Synaptic P-Rex1 signaling regulates hippocampal long-term depression and autism-like social behavior. Proc Natl Acad Sci U S A 112, E6964–6972.

Li, Z., Wu, K., Zou, Y., Gong, W., Wang, P., and Wang, H. (2022). PREX1 depletion ameliorates high-fat diet-induced non-alcoholic fatty liver disease in mice and mitigates palmitic acid-induced hepatocellular injury via suppressing the NF-kappaB signaling pathway. Toxicol Appl Pharmacol 448, 116074.

Lindsay, C.R., Lawn, S., Campbell, A.D., Faller, W.J., Rambow, F., Mort, R.L., Timpson, P., Li, A., Cammareri, P., Ridgway, R.A., et al. (2011). P-Rex1 is required for efficient melanoblast migration and melanoma metastasis. Nature Commun 2, 555.

Lorenzo-Martin, L.F., Rodriguez-Fdez, S., Fabbiano, S., Abad, A., Garcia-Macias, M.C., Dosil, M., Cuadrado, M., Robles-Valero, J., and Bustelo, X.R. (2020). Vav2 pharmaco-mimetic mice reveal the therapeutic value and caveats of the catalytic inactivation of a Rho exchange factor. Oncogene 39, 5098–5111.

Lucato, C.M., Halls, M.L., Ooms, L.M., Liu, H.J., Mitchell, C.A., Whisstock, J.C., and Ellisdon, A.M. (2015). The phosphatidylinositol (3,4,5)-trisphosphate-dependent Rac exchanger 1:Ras-related C3 botulinum toxin substrate 1 (P-Rex1:Rac1) complex reveals the basis of Rac1 activation in breast cancer cells. J Biol Chem 290, 20827-20840

Machin, P.A., Tsonou, E., Hornigold, D.C., and Welch, H.C.E. (2021). Rho family GTPases and Rho GEFs in glucose homeostasis. Cells 10.

Mauvais-Jarvis, F., Clegg, D.J., and Hevener, A.L. (2013). The role of estrogens in control of energy balance and glucose homeostasis. Endocr Rev 34, 309–338.

Menacho-Marquez, M., Nogueiras, R., Fabbiano, S., Sauzeau, V., Al-Massadi, O., Dieguez, C., and Bustelo, X.R. (2013). Chronic sympathoexcitation through loss of Vav3, a Rac1 activator, results in divergent effects on metabolic syndrome and obesity depending on diet. Cell Metab 18, 199–211.

Moller, L.L.V., Klip, A., and Sylow, L. (2019). Rho GTPases-emerging regulators of glucose homeostasis and Metabolic Health. Cells 8.

Montero, J.C., Seoane, S., and Pandiella, A. (2013). Phosphorylation of P-Rex1 at serine 1169 participates in IGF-1R signaling in breast cancer cells. Cellular signalling 25, 2281–2289.

Napari contributors. (2019). Napari: a multi-dimensional image viewer for python. doi:10.5281/zenodo.3555620.

Navale, A.M., and Paranjape, A.N. (2016). Glucose transporters: physiological and pathological roles. Biophys Rev 8, 5–9.

Osborn, O., Oh, D.Y., McNelis, J., Sanchez-Alavez, M., Talukdar, S., Lu, M., Li, P., Thiede, L., Morinaga, H., Kim, J.J., et al. (2012). G protein-coupled receptor 21 deletion improves insulin sensitivity in diet-induced obese mice. J Clin Invest 122, 2444–2453.

Pantarelli, C., Pan, D., Chetwynd, S., Stark, A.K., Hornigold, K., Machin, P., Crossland, L., Cleary, S.J., Baker, M.J., Hampson, E., et al. (2021). The GPCR adaptor protein Norbin suppresses the neutrophil-mediated immunity of mice to pneumococcal infection. Blood Adv 5, 3076–3091.

Reynolds, T.H., Dalton, A., Calzini, L., Tuluca, A., Hoyte, D., and Ives, S.J. (2019). The impact of age and sex on body composition and glucose sensitivity in C57BL/6J mice. Physiol Rep 7, e13995.

Riddy, D.M., Kammoun, H.L., Murphy, A.J., Bosnyak-Gladovic, S., De la Fuente Gonzalez, R., Merlin, J., Ziemann, M., Fabb, S., Pierce, T.L., Diepenhorst, N., et al. (2021). Deletion of GPR21 improves glucose homeostasis and inhibits the CCL2-CCR2 axis by divergent mechanisms. BMJ Open Diabetes Res Care 9.

Rodriguez-Fdez, S., Lorenzo-Martin, L.F., Fabbiano, S., Menacho-Marquez, M., Sauzeau, V., Dosil, M., and Bustelo, X.R. (2021). New functions of Vav family proteins in cardiovascular biology, skeletal muscle, and the nervous system. Biology 10, 857.

Rodriguez-Fdez, S., Lorenzo-Martin, L.F., Fernandez-Pisonero, I., Porteiro, B., Veyrat-Durebex, C., Beiroa, D., Al-Massadi, O., Abad, A., Dieguez, C., Coppari, R., et al. (2020). Vav2 catalysis-dependent pathways contribute to skeletal muscle growth and metabolic homeostasis. Nat Commun 11, 5808.

Rossman, K.L., Der, C.J., and Sondek, J. (2005). GEF means go: turning on Rho GTPases with guanine nucleotide-exchange factors. Nat Rev Mol Cell Biol 6, 167–180.

Shami, G.J, Samarska, I.V., Koek, G.H., Li, A., Palma, E., Chokshi, S., Wisse, E., and Braet, F. (2023). Giant mitochondria in human liver disease. Liver Intern 43, 2365–2378.

Thamilselvan, V., Gamage, S., Harajli, A., Chundru, S.A., and Kowluru, A. (2020). P-Rex1 mediates glucose-stimulated Rac1 activation and insulin secretion in pancreatic beta-cells. Cell Physiol Biochem 54, 1218–1230.

Thorens, B. (2015). GLUT2, glucose sensing and glucose homeostasis. Diabetologia 58, 221–232.

Wang, J., Pan, Z., Baribault, H., Chui, D., Gundel, C., and Veniant, M. (2016). GPR21 KO mice demonstrate no resistance to high fat diet induced obesity or improved glucose tolerance. F1000Res *5*, 136.

Welch, H.C. (2015). Regulation and function of P-Rex family Rac-GEFs. Small GTPases 6, 1–11.

Welch, H.C., Coadwell, W.J., Ellson, C.D., Ferguson, G.J., Andrews, S.R., Erdjument-Bromage, H., Tempst, P., Hawkins, P.T., and Stephens, L.R. (2002). P-Rex1, a PtdIns(3,4,5)P_3_- and Gβγ-regulated guanine-nucleotide exchange factor for Rac. Cell 108, 809-821.

Welch, H.C., Condliffe, A.M., Milne, L.J., Ferguson, G.J., Hill, K., Webb, L.M., Okkenhaug, K., Coadwell, W.J., Andrews, S.R., Thelen, M., et al. (2005). P-Rex1 regulates neutrophil function. Curr Biol 15, 1867–1873.

Xue, R., Lynes, M.D., Dreyfuss, J.M., Shamsi, F., Schulz, T.J., Zhang, H., Huang, T.L., Townsend, K.L., Li, Y., Takahashi, H., et al. (2015). Clonal analyses and gene profiling identify genetic biomarkers of the thermogenic potential of human brown and white preadipocytes. Nat Med 21, 760–768.

