## Supplemental figures and legends for "Prex1 controls glucose homeostasis by limiting glucose uptake and mitochondrial metabolism in liver through GEF-independent regulation of Gpr21"

Content:

Supplemental Figure Legends

Supplemental Figures 1-10

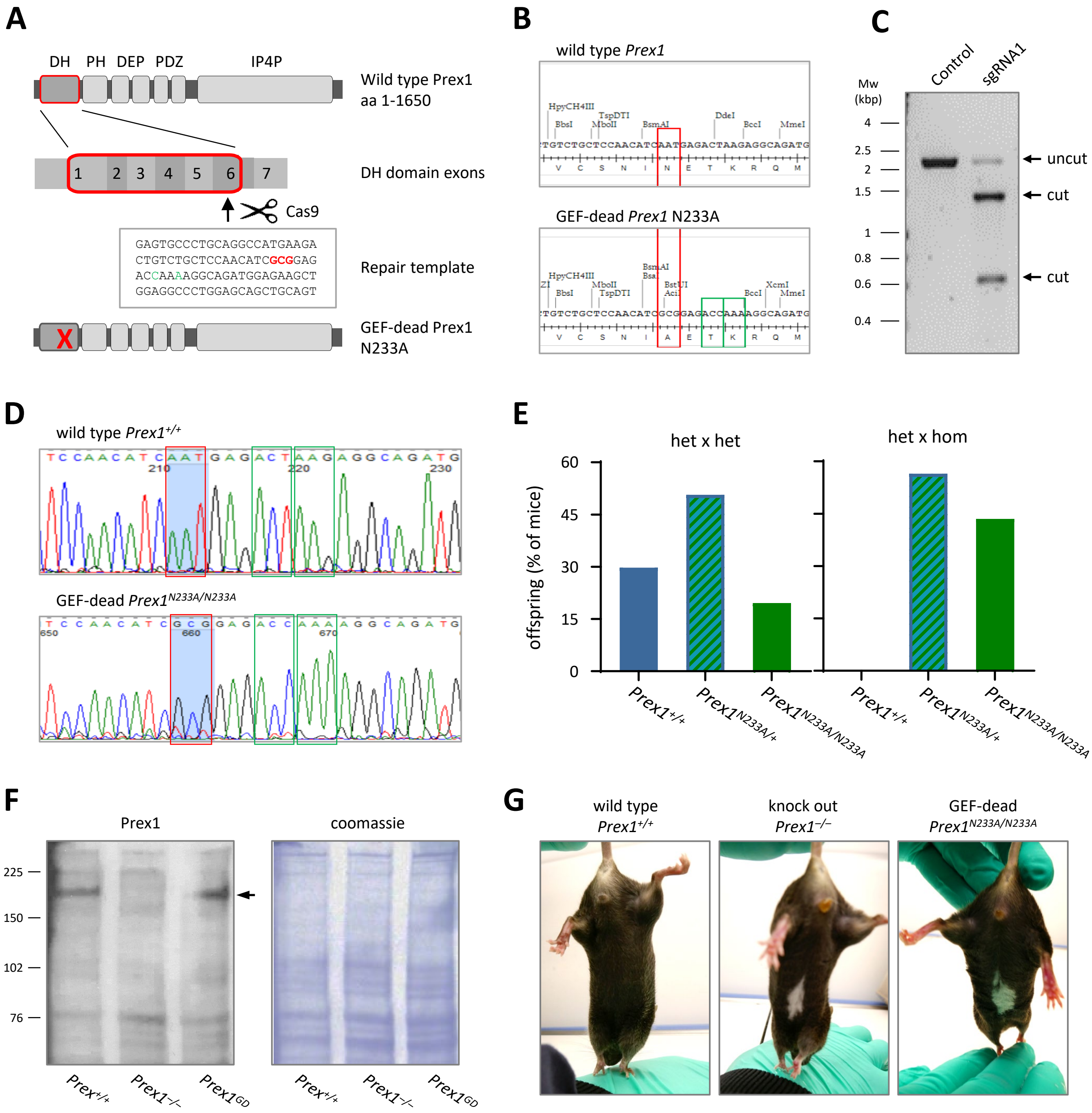


Supplemental Figure 1

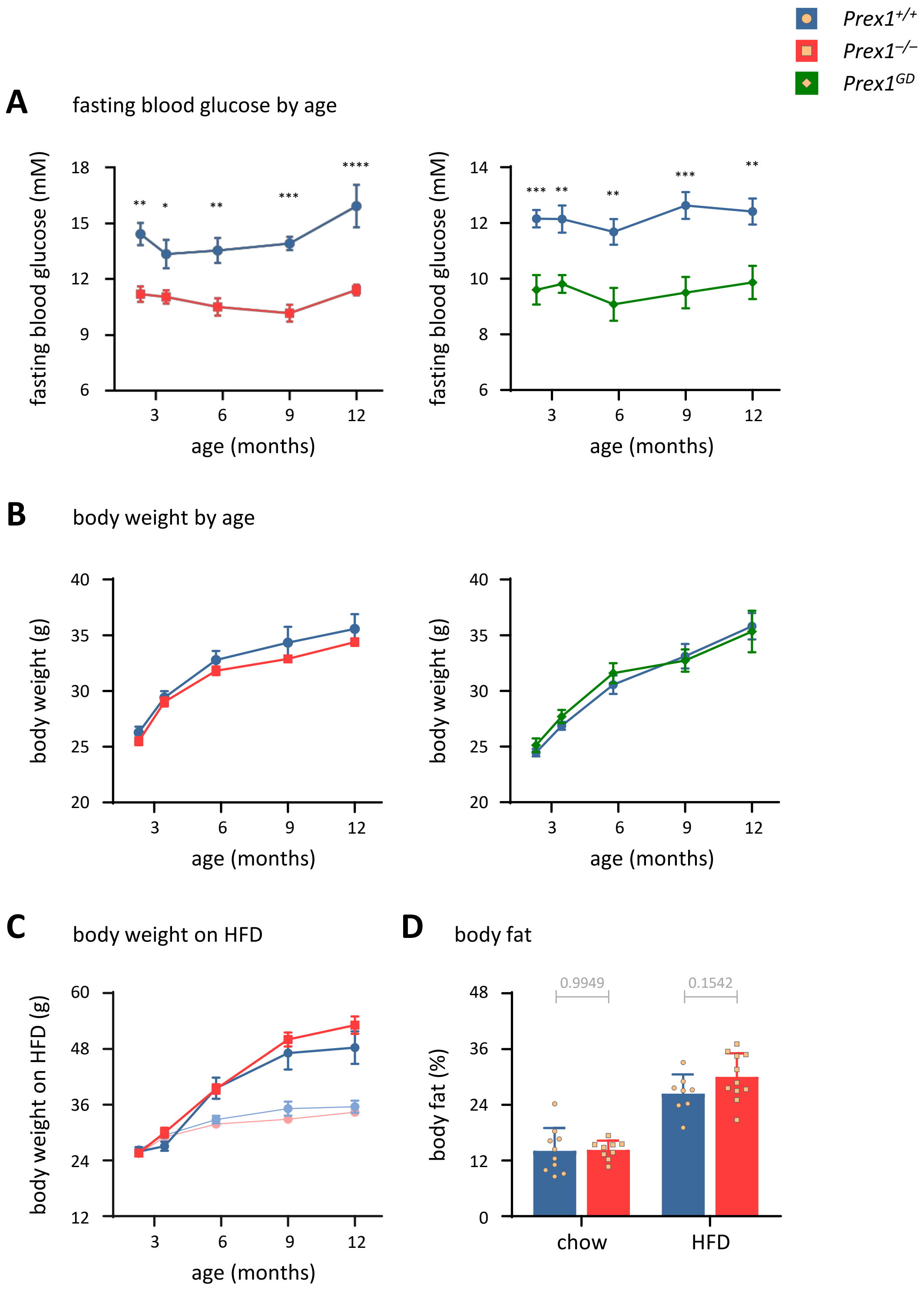


Supplemental Figure 2

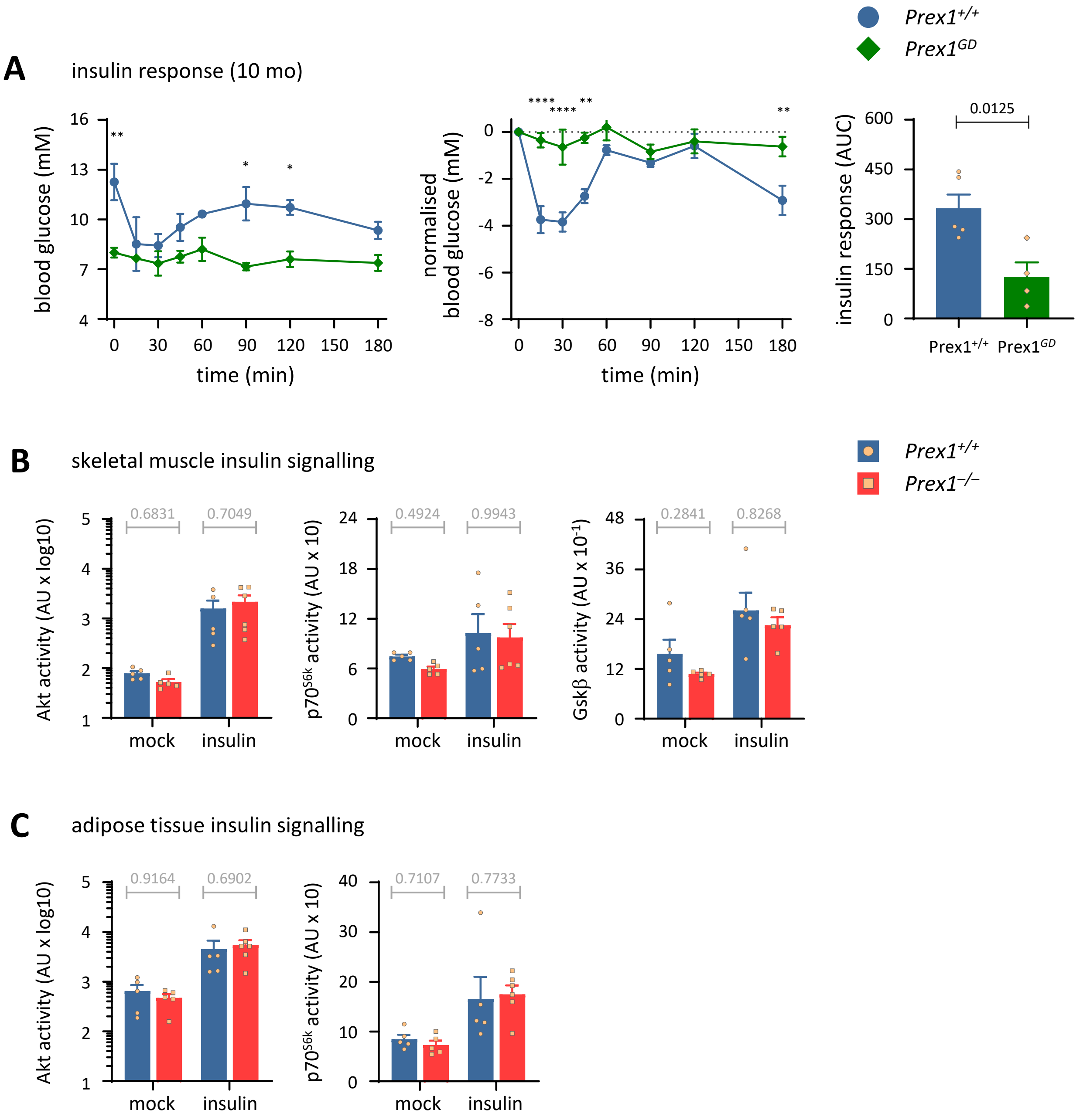


Supplemental Figure 3

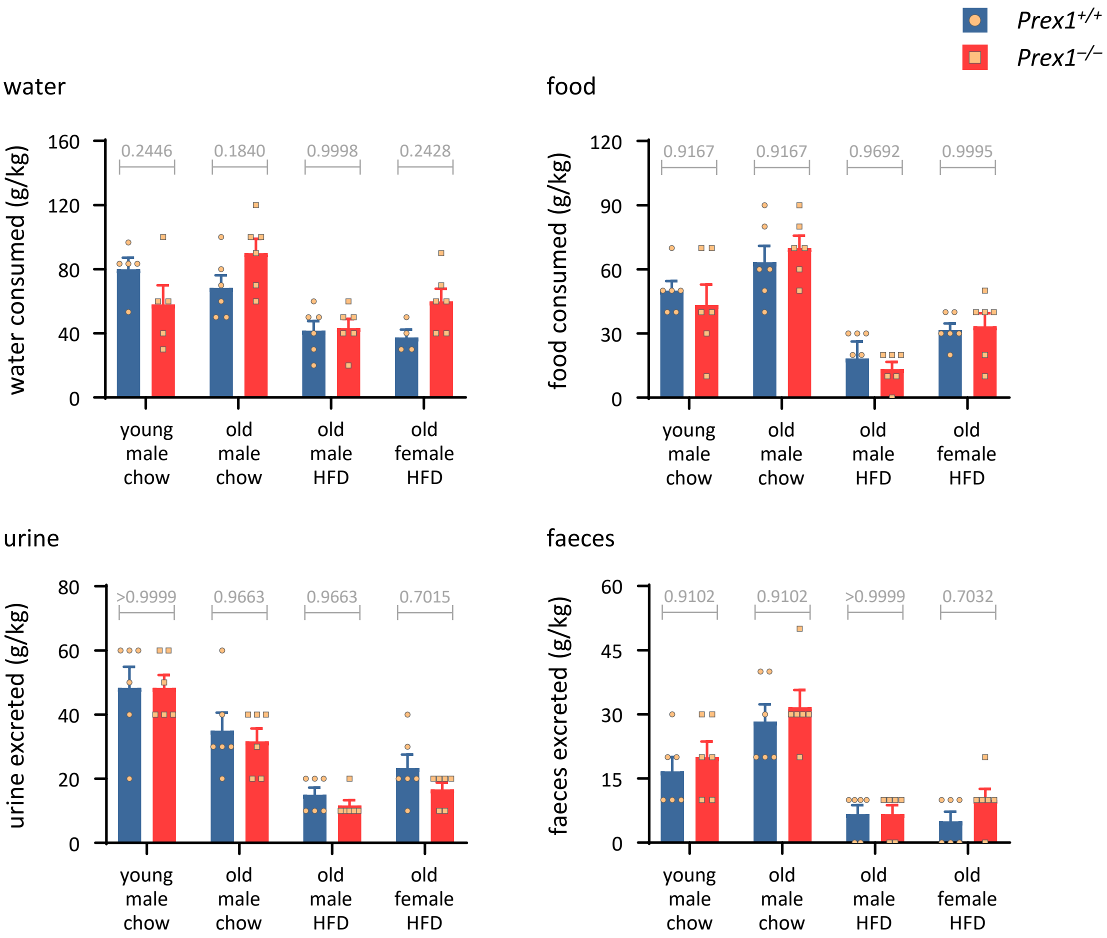


Supplemental Figure 4

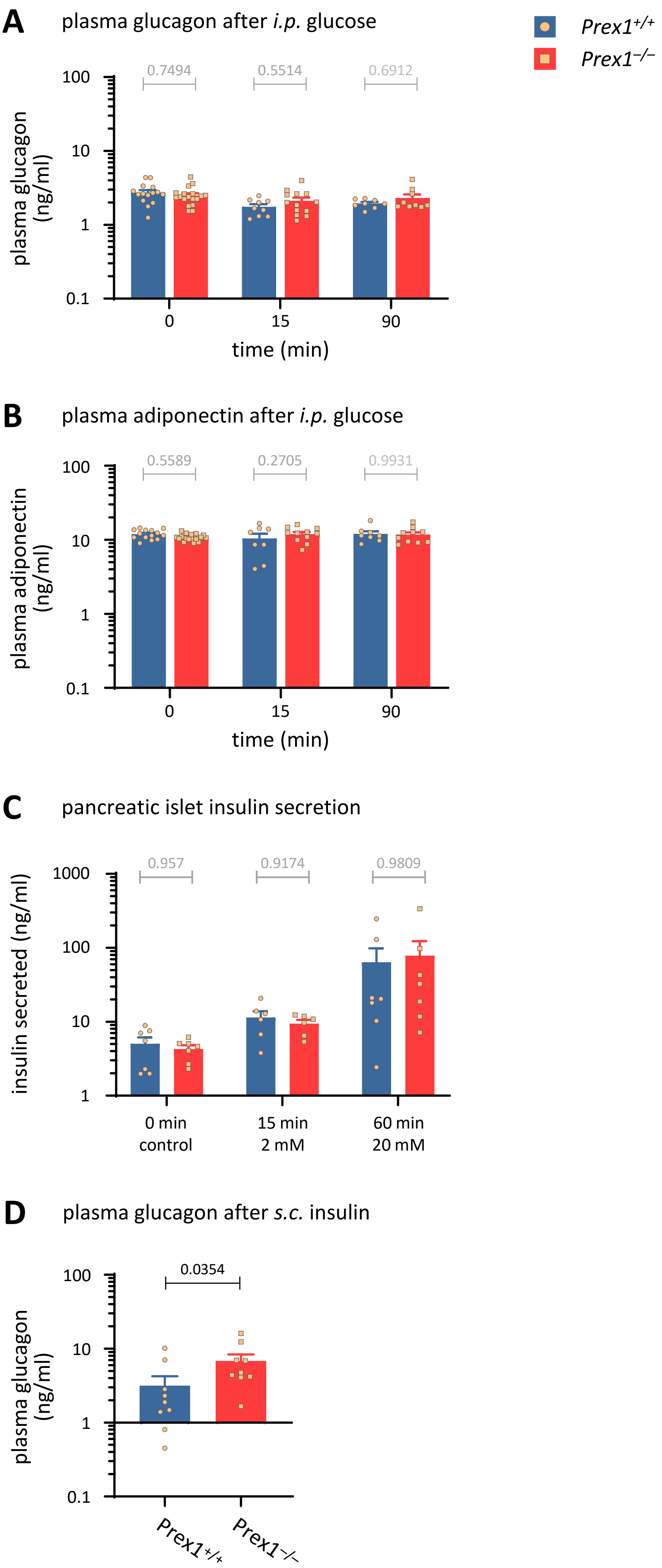


Supplemental Figure 5

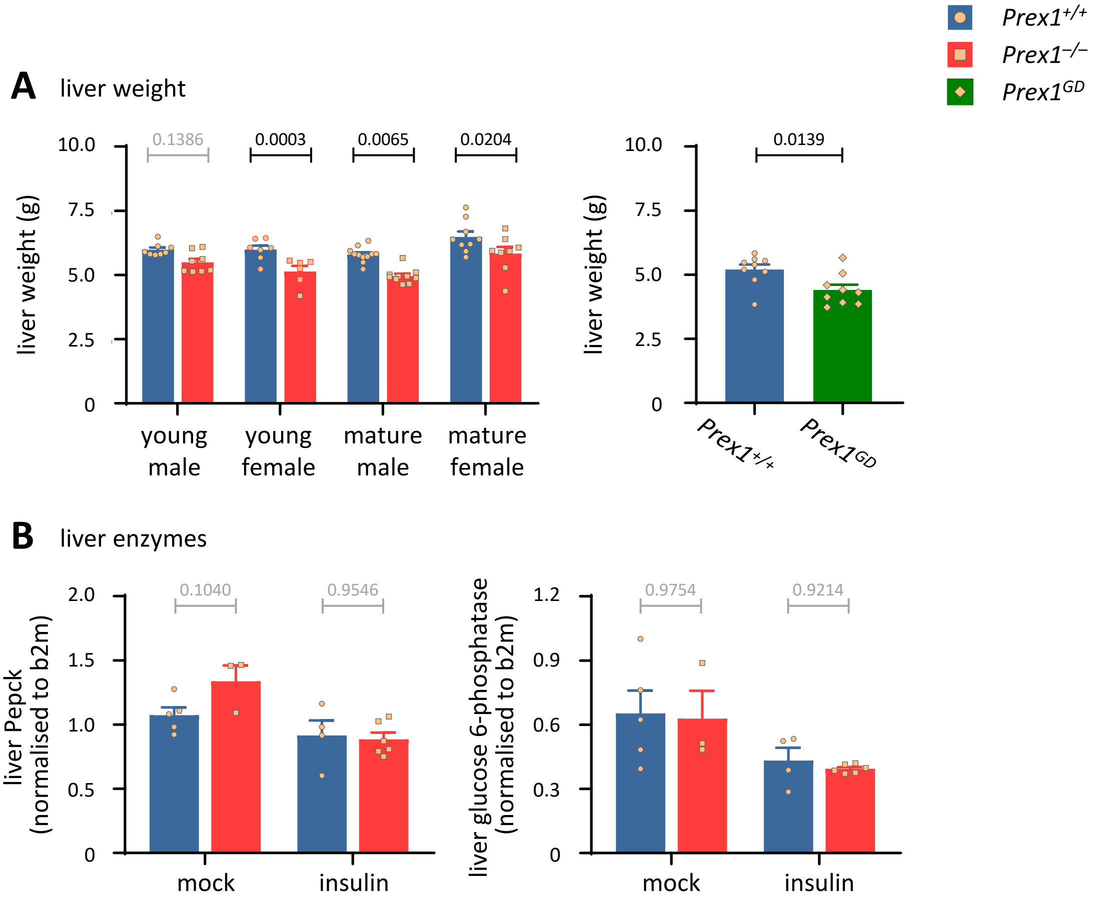


Supplemental Figure 6

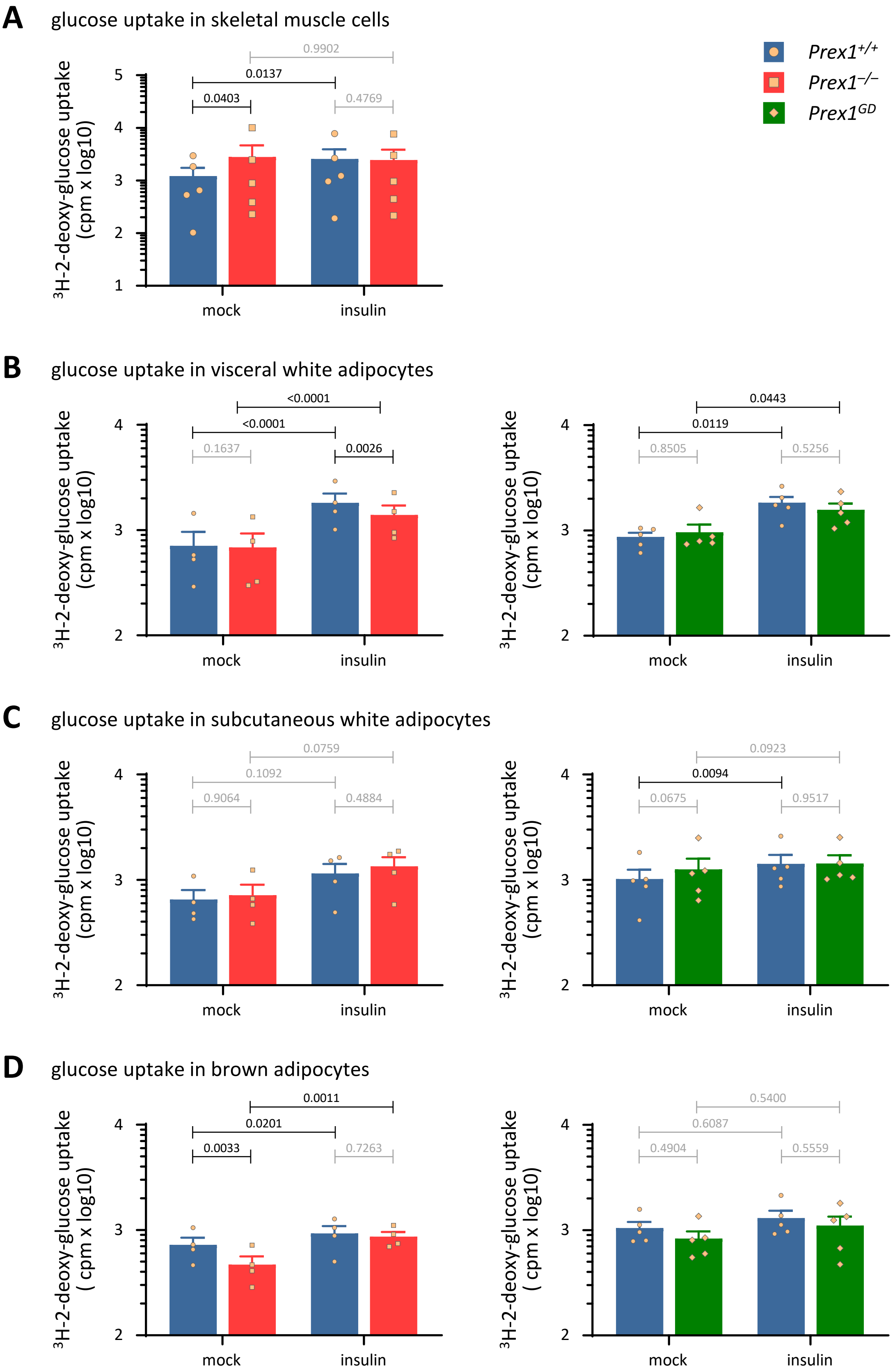


Supplemental Figure 7

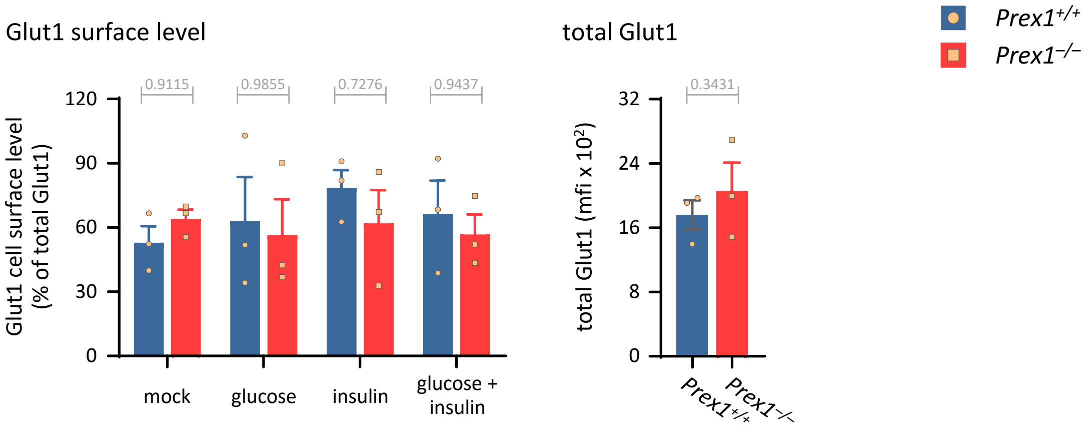


Supplemental Figure 8

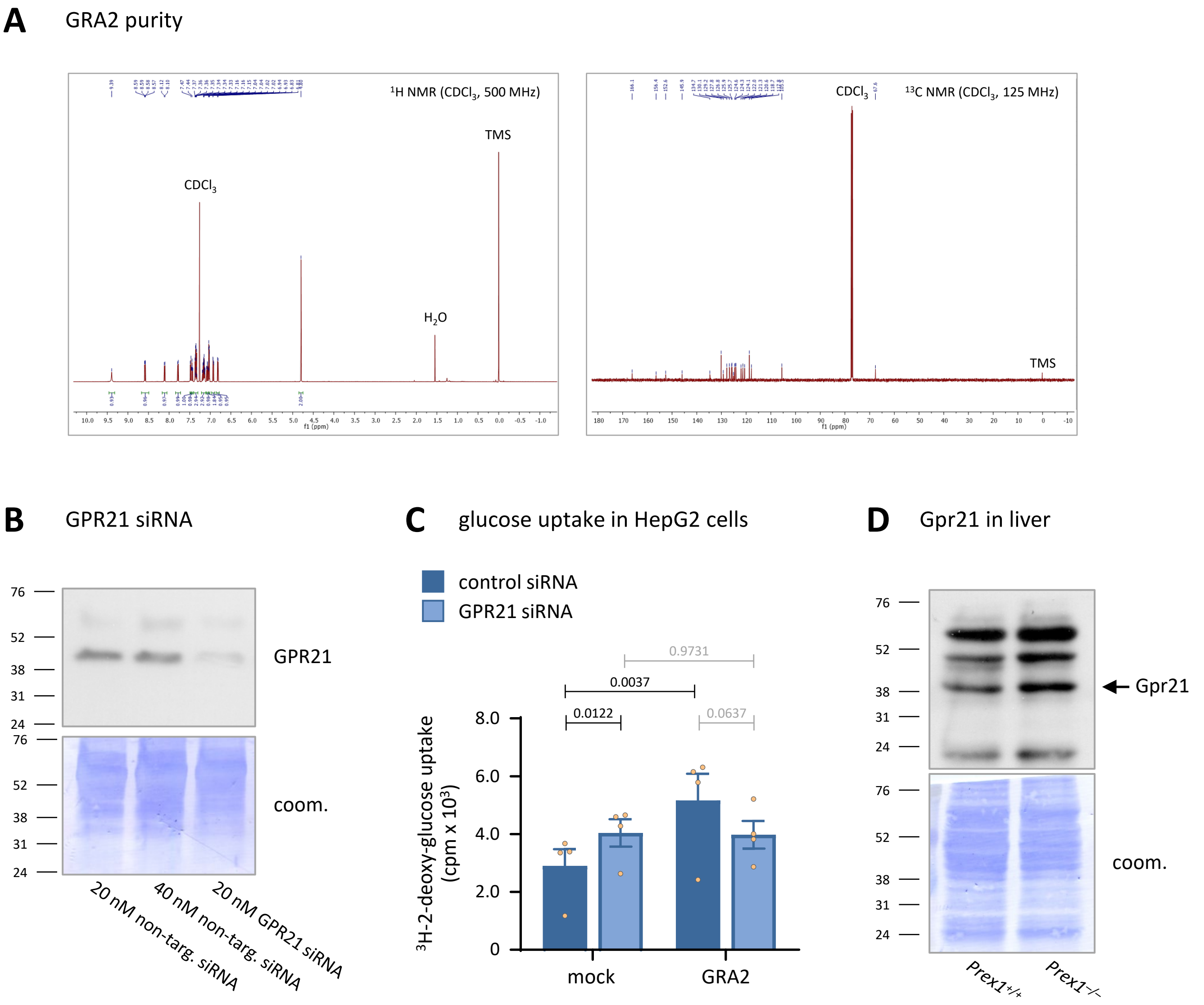


Supplemental Figure 9

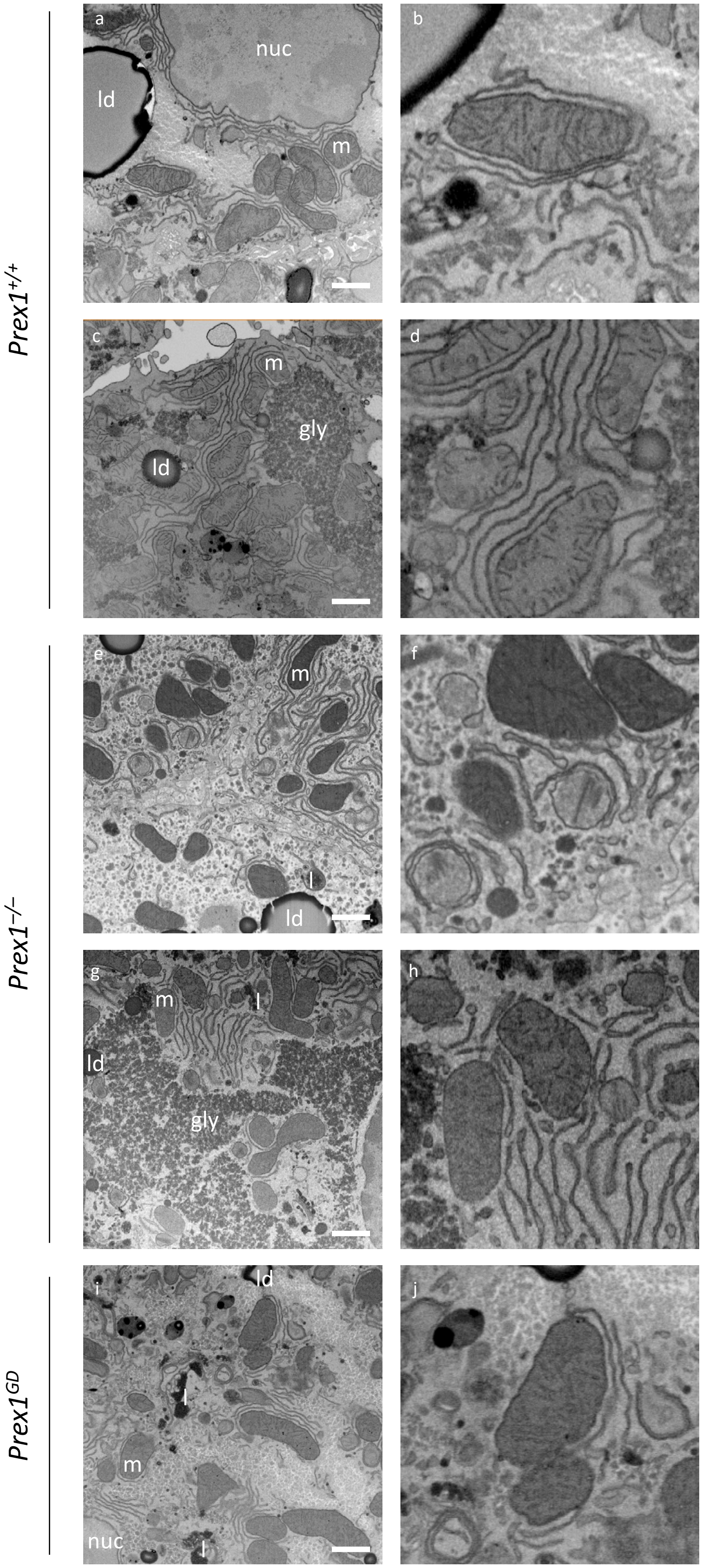


Supplemental Figure 10

### Supplemental Figure Legends

**Supplemental Figure 1. Generation of *Prex1*$^{GD}$ mice. (A)** Schematic of the targeting strategy. Cas9 nuclease was used to cut the target site in exon 6 in mouse *Prex1* which codes for N233 in the catalytic DH domain, and a repair template was used to introduce the N233A mutation and neighbouring silent mutations. **(B)** Restriction map of the wild type and targeted mouse genomic sequence, and translation. Red boxes show the nucleotide change which results in the N233A mutation. Green boxes highlight silent mutations introduced to alter restriction sites. **(C)** Assessment sgRNA efficiency *in vitro*. A PCR product of Prex1 mouse genomic sequence encompassing the target site in exon 6 was cleaved by recombinant Cas9 nuclease in the presence of sgRNA1. **(D)** Representative sequencing traces of a homozygous *Prex1*$^{GD}$ (*Prex1*$^{N233A/N233A}$) mouse compared to wild type. (B, D) Red boxes show the nucleotide change which results in the N233A mutation that renders *Prex1* catalytically inactive. Green boxes highlight silent mutations introduced to alter restriction sites. **(E)** % offspring of different genotypes from heterozygous (het) × het and from het x homozygous (hom) breeding pairs. Data are n = 306 pups from het × het crosses and 170 from het × hom crosses. **(F)** Normal expression level of Prex1$^{GD}$ in isolated neutrophils of *Prex1*$^{GD}$ mice. **(G)** White belly phenotype. *Prex1*$^{GD}$ mice have the white belly phenotype characteristic of *Prex1*$^{-/-}$ mice.

**Supplemental Figure 2. Prex1 raises fasting blood glucose levels throughout ageing, independently of its catalytic activity, without affecting body weight or body fat.** The fasting (6 h) blood glucose levels (A) and body weight (B) of the *Prex1*$^{+/+}$ (blue circles), *Prex1*$^{-/-}$ (red squares), and *Prex1*$^{GD}$ (green diamonds) mice in Figure 1A-C on chow diet are plotted as a function of age. Data are mean ± SEM of mice pooled from 2 independent cohorts of 4-5 mice per group, 10 *Prex1*$^{+/+}$ and 10 *Prex1*$^{-/-}$ mice (left), and 9 *Prex1*$^{+/+}$ and 10 *Prex1*$^{GD}$ mice (right). **(C)** Body weights of male Prex1$^{+/+}$ and Prex1$^{-/-}$ mice in Figure 1D on HFD as a function of age. Faint symbols with stippled lines show the body weights of mice from Figure 1A on chow diet, plotted for reference. Data are mean ± SEM of 8 *Prex1*$^{+/+}$ and 10 *Prex1*$^{-/-}$ mice pooled from 2 independent cohorts with 4-5 mice per group. Statistics in (A-C) are 2-way ANOVA with Sidak's multiple comparisons correction; stars denote significance between genotypes for the indicated ages. **(D)** The *Prex1*$^{+/+}$ and *Prex1*$^{-/-}$ mice in (A-C), on chow or HFD, were analysed by Echo MRI for % body fat. Data are mean ± SEM. Statistics are unpaired Student's *t*-test; p-values in grey are non-significant.

**Supplemental Figure 3. Prex1 is required for insulin sensitivity, through its Rac-GEF activity. (A)** Insulin sensitivity of *Prex1*$^{GD}$ mice. The insulin sensitivity of the *Prex1*$^{+/+}$ (blue circles) and *Prex1*$^{GD}$ (green diamonds) mice from Figure 1B and 2A was tested at the age of 10 months (43 weeks). Mice were fasted for 4 h, fasting blood glucose was measured, 0.375 IU/kg insulin *s.c.* injected, and blood glucose levels were tested at the indicated timepoints. Left-hand panel: blood glucose concentration, middle: response normalised to fasting blood glucose, right: integrated normalised response (AAC; beige dots show individual mice). Data are mean ± SEM of 5 *Prex1*$^{+/+}$ and 4 *Prex1*$^{GD}$ mice from one cohort. Statistics in time courses (A-C) are 2-way ANOVA with Sidak's multiple comparisons correction; stars denote significance between genotypes for the indicated time points. Statistics in the bar graphs are unpaired Student's *t*-test. **(B,C)** Mesoscale analysis of insulin signalling in skeletal muscle (B) and white adipose tissue (C). 10-week-old *Prex1*$^{+/+}$ (blue bars, circles) and *Prex1*$^{-/-}$ (red bars, squares) mice were fasted for 4 h, injected *s.c.* with 0.75 IU/kg insulin, or mock-treated, culled humanely after 15 min, tissues retrieved, homogenised, and active and total levels of Akt, p70$^{S6K}$, and GSK3β detected by Mesoscale analysis. Data are mean ± SEM of 5-6 mice per group, the same as in Figure 2C,D; beige symbols show individual mice. Statistics are 2-way ANOVA with Sidak's multiple comparisons correction on log-transformed raw data; p-values in grey are non-significant.

**Supplemental Figure 4. *Prex1*$^{-/-}$ mice perform normally in metabolic cages.** Young (12 weeks) and old (11 months) *Prex1*$^{+/+}$ (blue bars, circles) and *Prex1*$^{-/-}$ (red bars, squares) mice, male or female, and either on chow diet or on HFD from the age of 10 weeks, as indicated, were tested in metabolic cages on 3 subsequent nights with food and water provided *ad libitum*. Their water and food consumption, and their production of urine and faeces was measured after each night and normalised to body weight. Data are mean ± SEM mean for each night and cohort, with 2 independent cohorts per group (n=6). Young mice were 10 *Prex1*$^{+/+}$ and 10 *Prex1*$^{-/-}$ males, old mice were from the cohorts in Figures 1A, D, E. Statistics are 2-way ANOVA with Sidak's multiple comparisons correction; p-values in grey are non-significant.

**Supplemental Figure 5. Prex1 limits insulin-stimulated plasma glucagon but does not control glucose-stimulated insulin secretion from pancreatic islets. (A, B)** Glucose-stimulated plasma glucagon and adiponectin. Glucagon (A) and adiponectin (B) were measured in the plasma of 15 week-old *Prex1*$^{+/+}$ and *Prex1*$^{-/-}$males on chow diet fasted for 6 h (0 time), or fasted and challenged *i.p.* with 2 g/kg glucose for 15 or 90 min. Data are mean ± SEM of mice pooled from 3 independent cohorts (4 for 0'); beige symbols show individual mice, the same as in Figure 2B. Statistics are 2-way ANOVA on square root-transformed raw data with Sidak's multiple comparisons correction; p-values in black denote significant differences, p-values in grey are non-significant. **(C)** Glucose-stimulated secretion of insulin from pancreatic islets. Islets of Langerhans were isolated from the pancreas of 15-week-old *Prex1*$^{+/+}$ and *Prex1*$^{-/-}$ males on chow diet, and 10 islets were stimulated with 2 mM glucose for 15 min, then 20 mM glucose for a further 45 min. Insulin secreted into the medium was detected by ELISA. Data are mean ± SEM of 7 independent experiments; beige symbols show individual experiments. Statistics are 2-way ANOVA with Sidak's multiple comparisons correction; p-values in grey are non-significant. **(D)** Insulin-stimulated plasma glucagon. Glucagon was measured in the plasma of 14-week-old *Prex1*$^{+/+}$ and *Prex1*$^{-/-}$males on chow diet, after fasting for 4 h and challenge *s.c.* with 0.75 IU/kg insulin for 60 min. Data are mean ± SEM of 9 *Prex1*$^{+/+}$ and 9 *Prex1*$^{-/-}$males pooled from 2 independent cohorts of 4-5 per group; beige symbols show individual mice. Statistics are unpaired t-test on square root-transformed raw data.

**Supplemental Figure 6. Prex1-deficient mice have small livers but normal liver enzymes. (A)** Liver weights of young (10 weeks) and middle aged (males 24 weeks, females 30 weeks) *Prex1*$^{+/+}$ (blue bars, circles) and *Prex1*$^{-/-}$ (red bars, squares) mice (left) and of old *Prex1*$^{GD}$ (green bars, diamonds) mice (52 weeks, right) on chow diet. Data are mean ± SEM of 8-11 *Prex1*$^{+/+}$ and 6-11 *Prex1*$^{-/-}$ mice (right), and of 9 *Prex1*$^{+/+}$ and 9 *Prex1*$^{GD}$ mice (right), from two independent cohorts per group; beige symbols show individual mice. Statistics on the left are 3-way ANOVA with Sidak's multiple comparisons correction; on the right, unpaired t-test; p-values in black are significant, p-values in grey are non-significant. **(B)** Liver enzymes. *Pepck* and *G6p* in livers of *Prex1*$^{+/+}$ and *Prex1*$^{-/-}$ mice fasted for 4 h and *s.c.* injected with either DPBS (mock) or insulin at 0.75 IU/kg insulin for 15 min was quantified by qPCR. Data are mean ± SEM of 3-5 *Prex1*$^{+/+}$ and 4-6 *Prex1*$^{-/-}$ mice; beige symbols show individual mice. Statistics are 2-way ANOVA with Sidak's multiple comparisons correction.

**Supplemental Figure 7. Prex1 limits glucose uptake into skeletal muscle cells but does not control glucose uptake into adipose cells.** Glucose uptake was measured in **(A)** skeletal muscle cells, (**B)** mature visceral white, **(C)** subcutaneous white, and **(D)** brown adipose cells isolated from 15-week-old *Prex1*$^{+/+}$ (blue bars, circles), *Prex1*$^{-/-}$ (red bars, squares) and *Prex1*$^{GD}$ (green bars, diamonds) mice. Cells were stimulated with 100 nM insulin for 10 min at 37°C, or were mock-stimulated, followed by the addition of 50 µM 2-DOG, 0.25 µCi $H_3$-labelled 2-DOG for a further 60 min. Cells were washed, and glucose uptake was measured by scintillation counting. Data are mean ± SEM of 4-5 independent experiments; beige symbols show individual experiments. The different types of adipocytes were tested within the same experiments. Statistics are 2-way ANOVA with Sidak's multiple comparisons correction on log-transformed raw data; p-values in black are significant, p-values in grey are non-significant.

**Supplemental Figure 8. Prex1 does not affect Glut1 surface level in liver cells.** Liver cells from *Prex1*$^{+/+}$ (blue bars, circles) and *Prex1*$^{-/-}$ (red bars, squares) mice were stimulated with 5 mM glucose or 100 nM insulin for 10 min at 37°C, or were mock-stimulated, or stimulated with 100 nM insulin for 10 min and 5 mM glucose for another 30 min, stained with Glut1 antibody and analysed by flow cytometry. The mean fluorescence intensity (mfi) of Glut1 surface level is expressed as % of total Glut1. Data are mean ± SEM of 3 independent experiments; beige symbols show individual experiments. Statistics are 2-way ANOVA with Sidak's multiple comparisons correction. Right: Total Glut1 was measured in the same way except in permeabilised cells. Data are mean ± SEM of 3 independent experiments. Statistics are paired t-test.

**Supplemental Figure 9. Gpr21 antibody and GRA2 validation. (A)** Purity of GRA2. The purity of the GRA2 batch used was confirmed to be >95% by $^{1}H$ NMR (left) and $^{13}C$ NMR (right) spectroscopy, using deuterated chloroform ($CDCl_3$) as the solvent and a tetramethylsilane (TMS) internal standard. All signals other than those labelled are generated by GRA2. **(B)** HepG2 cells were transfected with non-targeting control siRNA or GPR21 siRNA for 72 h, and total lysates were western blotted with Gpr21 antibody. Coomassie staining was used as a loading control. Blots shown are representative of 2 independent experiments. **(C)** Glucose uptake in HepG2 cells. HepG2 cells transfected with non-targeting control siRNA or GPR21 siRNA for 72 h were treated with 30 mM GRA2 for 3 h at 37°C, or were mock-treated, followed by the addition of 50 µM 2-DOG, 0.25 µCi $^{3}H$-2-DOG for a further 60 min. Cells were washed, lysed and glucose uptake was measured by scintillation counting. Data are mean ± SEM of 4 independent experiments; beige symbols show individual experiments. Statistics are 2-way ANOVA with Sidak's multiple comparisons correction; p-values in black denote significant differences, p-values in grey are non-significant. **(D)** Total lysates of liver cells from *Prex1*$^{+/+}$ and *Prex1*$^{-/-}$ mice were western blotted with Gpr21 antibody. Coomassie staining was used as a loading control.

**Supplemental Figure 10. Prex1 prevents mitophagy in liver, independently of its catalytic Rac-GEF activity.** Ultrastructure of liver from 15-week-old male *Prex1*$^{+/+}$ (a-d), *Prex1*$^{-/-}$ (e-h), and *Prex1*$^{GD}$ (I, j) mice, observed in FIB-SEM XY slices from volumetric EM data sets. Mitochondria (m), nucleus (nuc), lipid droplets (ld), glycogen granules (gly) and lysosomes (l) are indicated. Images on the right are magnification regions of the left, highlighting mitochondria and mitophagy. Scale bars are 1 µm. Related to Figure 7A.
